# VASH1 phosphorylation regulates tubulin detyrosination via CNP/PRKG1α signaling in DRG neurons

**DOI:** 10.64898/2026.10.14.759141

**Authors:** Chadni Sanyal, Samar Ismail, Christophe Bosc, Eric Denarier, Jean-Marc Soleilhac, Béatrice Blot, Victoria Pakulska, Alexandra Kraut, Yohann Coute, Annie Andrieux, Lucie Carrier, Sacnicte Ramirez-Rios, Marie-Jo Moutin

## Abstract

Functional neuronal circuit formation requires precise regulation of axon growth and branching through dynamic cytoskeletal remodeling. Although microtubule detyrosination/re-tyrosination critically controls microtubule dynamics, how extracellular signals regulate this process remains unclear. Here, we identify a direct link between C-type natriuretic peptide, protein kinase cGMP-dependent 1α (CNP/cGMP/PRKG1α) signaling and microtubule detyrosination in dorsal root ganglion (DRG) neurons. CNP stimulation induces growth cone enlargement, increases tyrosinated tubulin within growth cones, and reduces detyrosinated tubulin along axons, effects abolished by PRKG1α inhibition. Pharmacological inhibition or genetic deletion of the detyrosinating enzyme VASH1 phenocopies CNP-induced growth cone remodeling. In vitro kinase assays identify VASH1 as a direct PRKG1α substrate and reveal a specific C-terminal serine phosphorylation site (serine 337) that suppresses VASH1-SVBP microtubule association and detyrosination activity. Together, these findings establish VASH1-SVBP as a key downstream effector linking extracellular guidance signaling to microtubule remodeling during DRG neuronal development.

**SUMMARY:** This study identifies VASH1-SVBP as a key downstream effector of the CNP/cGMP/PRKG1α signaling cascade, revealing how an extracellular cue regulates microtubule detyrosination and dynamics to shape growth cone morphology during dorsal root ganglion neuron development.

## INTRODUCTION

The establishment of correct neuronal circuits during nervous system development depends on the ability of growing axons to navigate through complex extracellular environments and form precise connections with their targets. Axon guidance and branching rely on the growth cone, a highly motile, actin- and microtubule-rich structure located at the distal tip of the developing axon, that interprets extracellular cues and converts them into cytoskeletal reorganization.

The growth cone is structurally compartmentalized into regions of specialized cytoskeletal composition. The peripheral domain mainly comprises of dynamic microtubules and F-actin in the form of lamellipodia and filopodia, while the central domain is dominated by stable microtubules that extend from the axon shaft. Growth cone guidance and protrusion are coordinated through dynamic interactions between actin and microtubules (Dent and Gertler, 2003; Lowery and Van Vactor, 2009). Microtubules provide mechanical stability and intracellular transport rails, and their dynamics are tightly regulated by numerous microtubule-associated proteins (MAPs) and post-translational modifications (PTMs). One of the best-characterized PTMs is the reversible detyrosination/re-tyrosination cycle of α-tubulin, which exerts a profound influence on microtubule dynamics, motor proteins binding and intracellular cargoes transport, and therefore neuron development (Janke and Magiera, 2020; Moutin et al., 2021; Sanyal et al., 2021).

The tubulin detyrosination/tyrosination cycle is catalyzed by opposing enzymatic systems. The removal of the C-terminal tyrosine present on most α-tubulin (detyrosination) is carried out by VASH-SVBP complexes (vasohibin 1 or 2 in complex with the small vasohibin-binding protein (Aillaud et al., 2017; Nieuwenhuis et al., 2017; Ramirez-Rios et al., 2023)) and MATCAP (microtubule associated tyrosine carboxypeptidase, also called tubulin metallocarboxypeptidase TMCP1 (Landskron et al., 2022; Nicot et al., 2023)). The tubulin tyrosine ligase (TTL) re-adds the tyrosine residue to α-tubulin (Ersfeld et al., 1993). This enzymatic balance defines spatially separated populations of microtubules with, in neurons, only dynamic, tyrosinated microtubules near the growth cone front and both dynamic, tyrosinated microtubules and stable, detyrosinated microtubules extending into axon shafts (Ahmad et al., 1993; Wehland and Weber, 1987). Although detyrosination/re-tyrosination of tubulin is known to be crucial for neuronal differentiation and axonal architecture (Erck et al., 2005; Pagnamenta et al., 2019; Peris et al., 2009), and TTL required for proper morphology and functionality of the growth cone (Marcos et al., 2009), how the enzymes of the cycle are regulated in developing neurons is unknown.

Sensory dorsal root ganglia (DRG) neurons provide an excellent model to study how the cytoskeleton could be controlled by extracellular guidance molecules. These neurons have a stem axon which bifurcates into a peripheral and a central axon to reach defined anatomical points (Nascimento et al., 2018). One of the signalling pathways involved in DRG axon development is mediated by C-type natriuretic peptide (CNP), a member of the peptide family that also consists of A- and B-type natriuretic peptides. Unlike its endocrine counterparts, CNP acts locally in sensory neurons to regulate axon outgrowth, bifurcation and guidance (Kishimoto et al., 2008; Zhao and Ma, 2009). Upon binding to its receptor NPR2 (natriuretic peptide receptor 2 or B), a membrane-bound guanylyl cyclase, CNP stimulates the intracellular generation of cyclic guanosine monophosphate (cGMP), activating protein kinase cGMP-dependent type 1 (PRKG1, also named cGK1) (Dumoulin et al., 2018b). Of the two PRKG1 isoforms, PRKG1α predominates in sensory neurons, and loss of either CNP or PRKG1α leads to profound DRG axon bifurcation defects and aberrant sensory wiring (Dumoulin et al., 2018b; Schmidt et al., 2002; Zhao and Ma, 2009). These findings identify the CNP/cGMP/PRKG1α axis as a critical regulator of axon guidance in sensory neurons (Dumoulin et al., 2021). Given the essential role of microtubule dynamics and PTMs such as detyrosination/re-tyrosination in shaping growth cone structure, an intriguing possibility is that CNP/cGMP/PRKG1α signalling may regulate the microtubule-modifying enzymes involved in this tubulin modification cycle, thereby linking extracellular signalling to intracellular cytoskeletal remodelling. We thus hypothesized that PRKG1α signalling may directly interface with the detyrosination/re-tyrosination machinery. To test this hypothesis, embryonic mouse DRG explants and cultures, pharmacological modulators, and biochemical assays were used to dissect the signalling pathway from CNP to the cytoskeletal level. We showed that CNP, via PRKG1α, promotes DRG growth cone enlargement accompanied with significantly increased tyrosinated tubulin within the growth cone and decreased detyrosinated tubulin along the axon. EpoY, the irreversible inhibitor of VASH-SVBP enzymes, phenocopied the CNP-induced growth cone enlargement and tubulin modifications pattern, while genetic deletion of VASH1 abolished the effect of CNP. Furthermore, PRKG1α phosphorylates in vitro the serine residue 337 (S337) of recombinant VASH1 within its C-terminal domain, and mutating this residue to glutamate (S337E phosphomimic) notably reduced the binding to microtubules and activity of the detyrosinase. Furthermore, using a newly developed antibody, we evidenced the presence of the S337-phosphorylated form of VASH1 in DRG neurons, and revealed its accumulation under CNP treatment. Overall, our results demonstrate that VASH1 acts as an essential effector downstream of the CNP/cGMP/PRKG1α pathway, linking extracellular signalling to intracellular cytoskeletal remodeling in developing neuron.

## RESULTS

### CNP mediates changes in tubulin tyrosination balance and growth cone enlargement via PRKG1 in DRG neurons

We first examined the effect of CNP on the growth cone area of cultured DRG explants obtained from embryonic day 13 (E13) wild-type (WT) mouse embryos (E13, Figure 1A). Growth cones were visualized with phalloidin, a naturally occurring toxin that targets F-actin. We observed that a 1 h treatment with CNP (500 nM) increased the average growth cone area of DRG neurons by 73% (Figure 1B, C), confirming previous findings (Dumoulin, Dagane, et al., 2018). As tyrosination of tubulin was shown to be important for proper growth cone morphogenesis and organization during development (Marcos et al., 2009), we simultaneously examined tyrosinated (Tyr) and detyrosinated (dTyr) tubulin in DRG neurons using specific antibodies against these two species of tubulin. Tyr tubulin beared by dynamic microtubules was clearly detected within the growth cone structure while dTyr tubulin was enriched along stable axonal microtubules (Figures 1D), which is consistent with previous studies (Baas and Black, 1990; Brown et al., 1993)(for review see (Moutin et al., 2021)). The growth cone enlargement induced by CNP was accompanied by a significant increase (55%) in the Tyr tubulin levels within the growth cone structure (Figure 1E).

**Figure 1.**
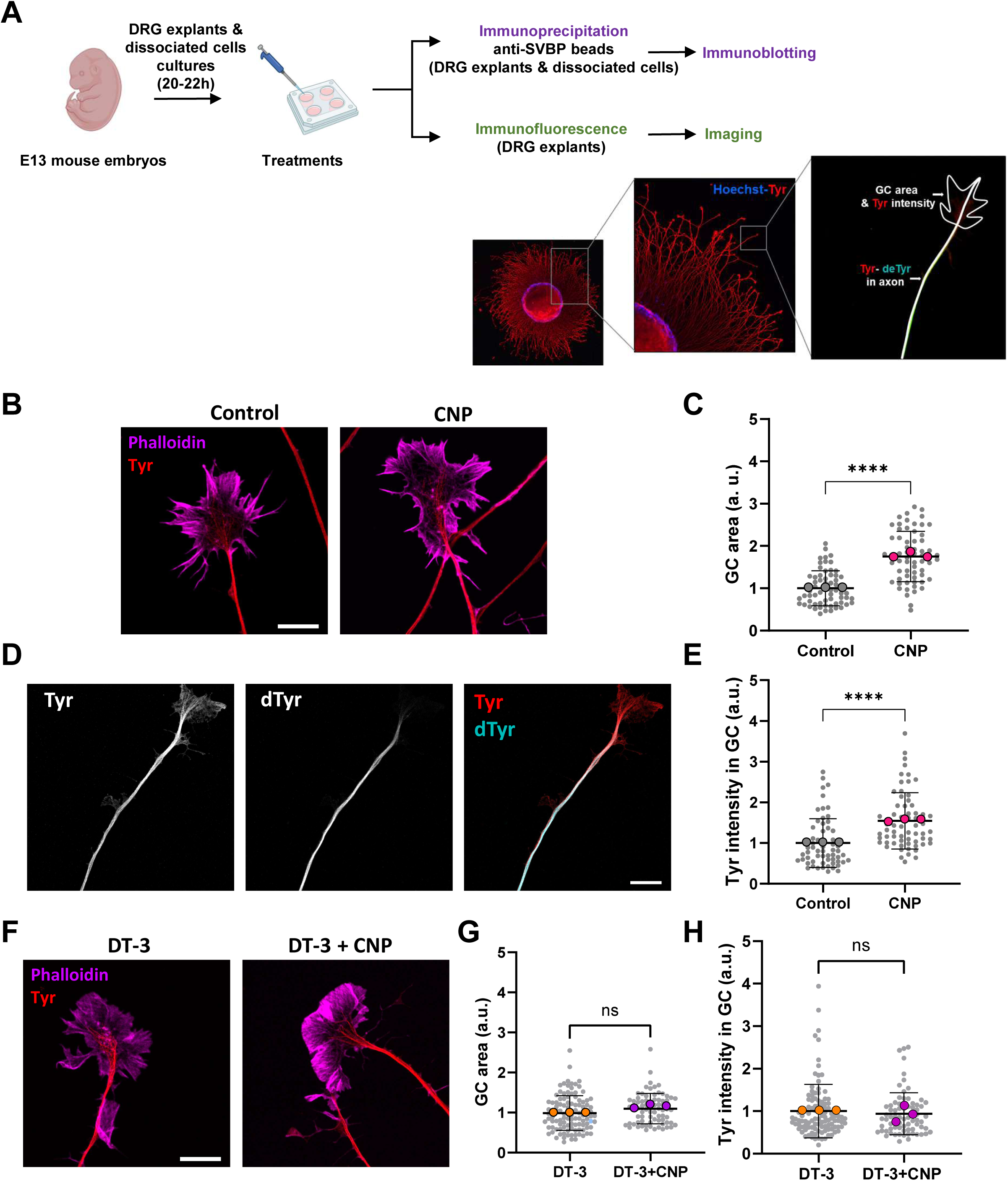
CNP impacts growth cone area and tyrosination levels of DRG neurons through a PRKG1 signalling. (**A**) Scheme of the protocols employed. DRG explants or dissociated neurons from embryonic day 13 (E13) mice were grown for 20-22 h before varying treatments. They were then employed for immunofluorescence or immunoprecipitation experiments followed by immunoblots. Representative immunofluorescence images of an embryonic DRG stained for tyrosinated tubulin is presented together with a scheme of the tubulin species (tyrosinated = Tyr; detyrosinated = dTyr) analyzed in the neuronal compartments. DRG explants were pretreated for 2 h with either vehicle (DMSO, control) or a PRKG1α inhibitor (DT-3, 150 nM), followed by treatment with or without CNP (500 nM, 1 h), as indicated on the Figure. **(B, F)** Representative confocal images of growth cones stained for tyrosinated α-tubulin (Tyr, red) and F-actin (phalloidin, light magenta). Scale bar, 10 µm. Quantifications of area (**C, G**) and tyrosinated tubulin intensity (**E, H**) of growth cones. n/N: 61/3 (Control), 64/3 (CNP), 105/3 (DT-3), 74/3 (DT-3+CNP), with n indicating the number of growth cones and N independent experiments. Data are expressed as mean ± SD and as fold change over control (**C, E**) or DT-3 (**G, H**). Large data points represent the mean value of each experiment. Statistical significance was determined using Mann-Whitney test. ****p < 0.0001, ns p > 0.05. a.u., arbitrary unit. (**D**) Representative confocal images of axons stained for tyrosinated (Tyr, red) and detyrosinated α-tubulin (dTyr, turquoise). Scale bar, 10 µm.

Based on an earlier study (Dumoulin et al., 2018a), the next vital target to confirm downstream of CNP was the implicated kinase, PRKG1. To address this, we used the cell permeable inhibitory peptide (DT-3, 150 nM) and examined the effect of CNP following kinase inhibition in WT DRG explants. As expected, the presence of DT-3 prevented the growth cone enlargement induced by CNP (Figure 1F, G). Interestingly, in the presence of the kinase inhibitor, Tyr tubulin levels did not differ in growth cone of DRG explants treated with CNP or vehicle (Figure 1H).

Our data therefore validate the role of PRKG1 downstream of CNP on the growth cone morphology of DRG neurons, together with revealing an impact of this pathway on Tyr tubulin levels within their growth cones. Thus, we wondered whether a tubulin detyrosination enzyme could be the target of the cGMP-dependent kinase to explain the changes in tubulin in these neurons.

### VASH1-SVBP is a downstream mediator of CNP-induced growth cone enlargement

The VASH1/2-SVBP complexes, formed by vasohibin (VASH1 or VASH2) and small vasohibin-binding protein (SVBP), are the major tubulin detyrosinating enzymes in cells and tissues (Aillaud et al., 2017; Nieuwenhuis et al., 2017). These enzymes, which exhibit high catalytic activity, are generally expressed at low levels (Aillaud et al., 2017; Ramirez-Rios et al., 2023). To detect their expression in DRG neurons, we used an immuno-precipitation method employing a custom-made anti-SVBP antibody (Figure 1A, see also Materials and Methods). We revealed the presence of VASH1, but not VASH2, in extracts and explants of DRG (Figure 2A).

**Figure 2.**
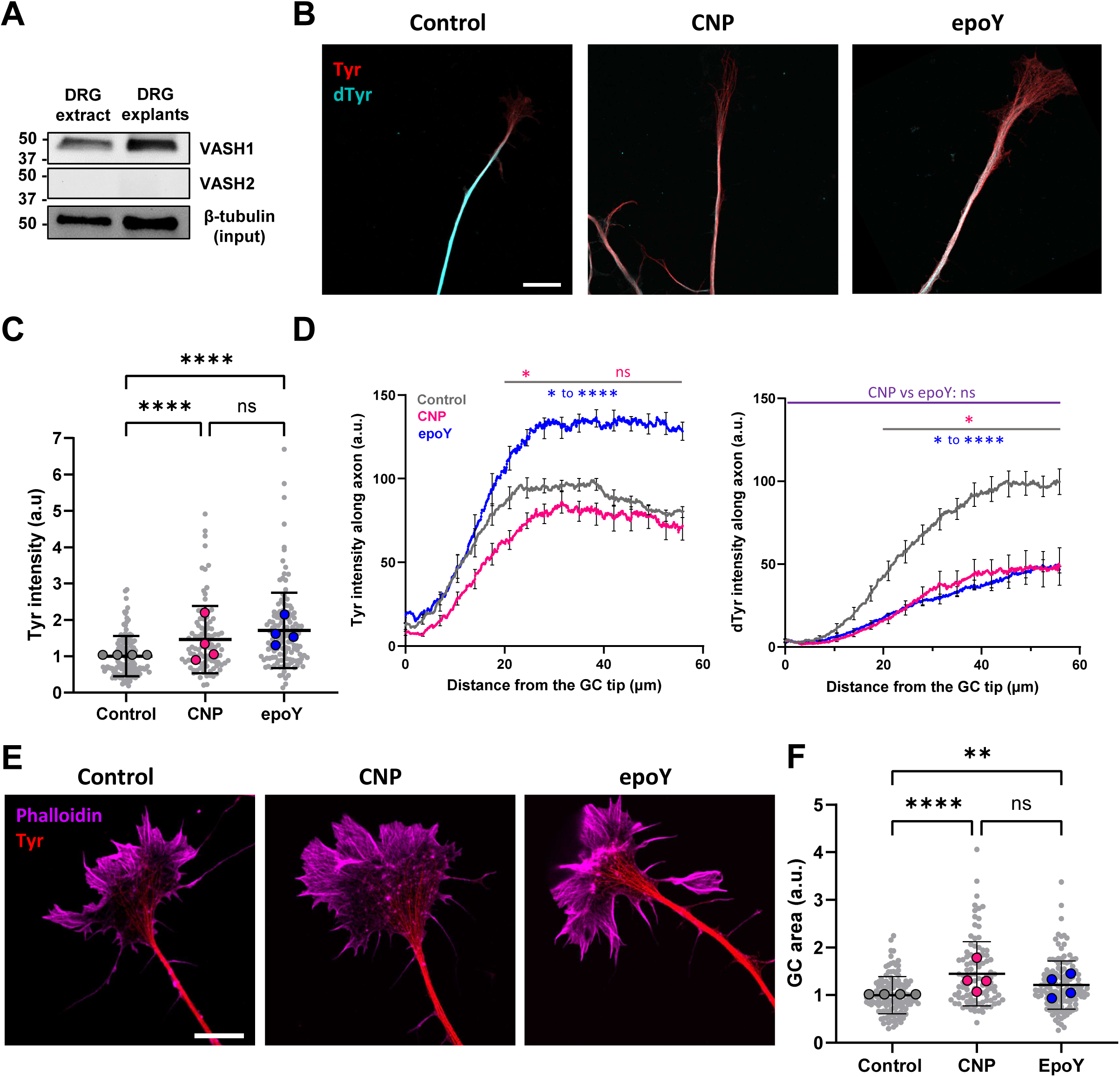
The VASH detyrosinating enzymes inhibitor, epoY, induces CNP-like changes in the growth cone and axon of DRGs. (**A**) Detection of VASH1 protein by immunoblot in DRG extract and cultured explants after immunoprecipitation using an anti-SVBP antibody. β-tubulin signal confirms similar protein content in the immunoprecipitation inputs. (**B-F**) E13 mouse DRG explants were treated with vehicle (DMSO control), CNP (500 nM, 1 h) or epoY (50 µM, 3 h), as indicated in the panels. **(B)** Representative confocal images of axons stained for tyrosinated (Tyr, red) and detyrosinated α-tubulin (dTyr, turquoise). Scale bar, 10 µm. See also Figure S2A. (**C**) Quantification of tyrosinated (Tyr) α-tubulin levels in growth cone. n/N: 126/4 (Control), 101/4 (CNP), 134/4 (epoY), with n indicating number of growth cones and N independent experiment. Data are expressed as mean ± SD and as fold change over control. Large data points represent the mean value of each experiment. Statistical significance was determined using Kruskal-Wallis test. ****p < 0.0001, ***p < 0.001, ns p > 0.05. (**D**) Quantification of tyrosinated (Tyr) and detyrosinated (dTyr) α-tubulin along the axons (“0” being the tip of the growth cone). n/N: 115/8 (Control), 87/8 (CNP), 85/6 (epoY) with n indicating number of axons and N number of slides (2 per experiment). Data are expressed as mean ± SEM (using N) and relative to control. Statistical significance was determined using two-way ANOVA and Tukey’s multiple comparison test. ***p < 0.001, **p < 0.01, *p < 0.05, ns p > 0.05. CNP and epoY vs Control are represented in pink and blue, respectively; CNP vs epoY in purple (**E**) Representative confocal images of growth cones stained for tyrosinated α-tubulin (Tyr, red) and F-actin (Phalloidin, magenta). Scale bar, 10 µm. (**F**) Quantification of growth cone areas. Data are expressed as mean ± SD and as fold change over control. N, n and statistics are same as in (**C**). a.u.: arbitrary unit.

We then tested whether a VASH-SVBP activity could be implicated in the CNP-induced morphological change of growth cone by using epoY, a known irreversible inhibitor of the enzyme (Aillaud et al., 2017). We compared the effect of the drug to that of CNP on DRG neurons. EpoY (50 µM, 4 h) increased overall Tyr tubulin within the growth cone, similar to CNP (Figures 2B, 2C, S2A; mean increases of 40% for CNP and 65% for epoY). We also analysed Tyr and dTyr tubulin changes along the axon following CNP and epoY treatments by tracing a line from the growth cone front and measuring fluorescence intensities along this line (scheme Figure 1A). The levels of both Tyr and dTyr tubulin progressively increased from the growth cone tip and along the axon, where microtubules are known to accumulate as compacted structures (Figures 2B, 2D, S2A). CNP treatment significantly decreased dTyr tubulin levels all along DRG axons while not significantly changing axonal Tyr tubulin levels, although a downward trend was observed (Figures 2D, S2A). Thus, CNP reduced tubulin detyrosination in axon and promoted redistribution of Tyr tubulin toward the growth cone. These CNP-induced changes were not observed when preincubating with the PRKG1 inhibitor DT-3 (Figure S2B, C). EpoY drastically reduced dTyr tubulin levels along the axon (Figures 2D, S2A), resembling CNP effect. Unlike CNP, epoY also significantly elevated Tyr tubulin intensity along the axon (Figures 2D, S2A). This difference may result from the irreversible and global action of epoY, leading to an accumulation of Tyr tubulin pools throughout the axon, whereas CNP, acting through PRKG1, may exert a more localized effect near the growth cone. Remarkably, epoY treatment also led to a growth cone enlargement (23%, Figure 2E, F), although less pronounced than that observed under CNP treatment.

The similar profiles observed with both epoY and CNP treatments suggest that VASH1-SVBP, that is present in DRGs, is a downstream effector of the CNP/cGMP/PRKG1 pathway. To confirm this hypothesis, we utilised DRG explants isolated from E13 *Vash1* knockout (KO) mouse models developed by the CRISPR/Cas9 technology (see Material and Methods and Figure S3A). Our initial observations showed that the mean growth cone area was ∼38% higher in DRGs from *Vash1* KO than WT (Figure 3A, B). Similarly, the mean levels of Tyr tubulin within DRGs growth cones were 2.6-fold higher in *Vash1* KO than in WT (Figure 3C, D). Next, we examined the effect of CNP on the DRGs explants from *Vash1* KO embryos and found that the absence of the enzyme abolished the CNP-mediated growth cone enlargement (Figure 3E, F). Notably, no changes in Tyr tubulin levels within the growth cone were observed upon CNP treatment in *Vash1* KO DRGs (Figure 3G, H). Finally, in contrast to WT DRGs, CNP did not alter the tyrosinated/detyrosinated tubulin balance along the axons of *Vash1* KO DRGs (Figure S3B).

**Figure 3.**
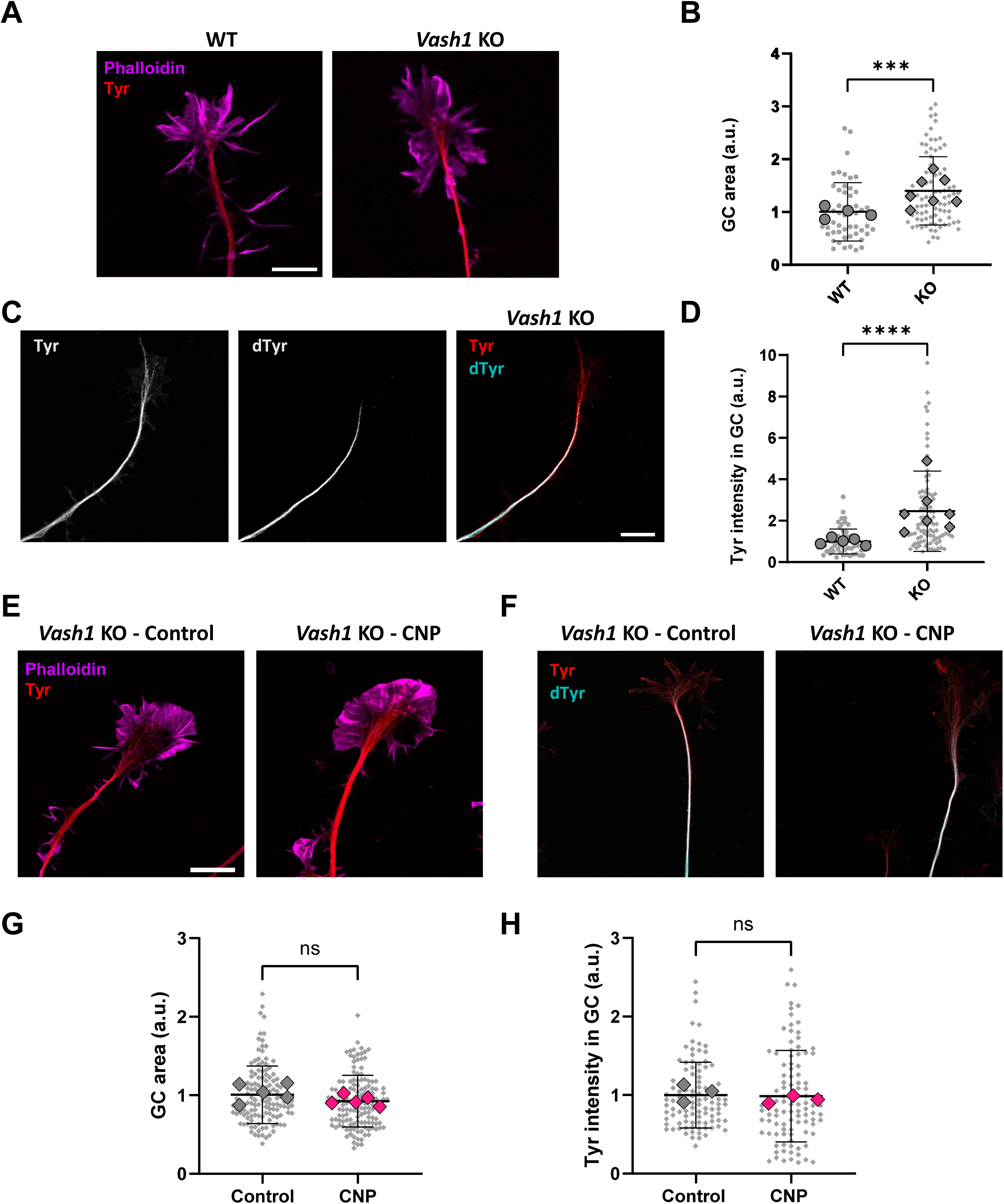
The loss of VASH1 hinders CNP-induced enlargement of DRG growth cones. (**A-D**) Comparison of DRG explants from wild-type (WT) and *Vash1* knockout (KO) E13 mice. Representative confocal images of **(A**) growth cones stained for tyrosinated α-tubulin (Tyr, red) and F-actin (Phalloidin, magenta) and (**C**) axons stained for tyrosinated (Tyr, red) and detyrosinated α-tubulin (dTyr, turquoise). Scale bar, 10 µm. Comparison of area **(B)** and tyrosinated (Tyr) α-tubulin signal **(D)** of the DRG growth cones from WT and *Vash1* KO mice. In each experiment, the signal values for the KO were relative to those of the WT. n/N: in (**B**), 51/4 (WT), 87/7 (*Vash1* KO) and in (**D**), 58/4 (WT), 93/7 (*Vash1* KO), with n indicating number of growth cones and N number of mice embryos. Data are expressed as mean ± SD and as fold change over WT. Statistical significance was determined using Mann-Witney test. ****p < 0.0001, ***p < 0.001, ns p > 0.05. a.u.: arbitrary unit. (**E-H**) DRG explants from *Vash1* KO E13 mice were treated with vehicle or CNP (500 nM, 1 h). Representative confocal images of **(E)** growth cones stained for tyrosinated α-tubulin (Tyr, red) and F-actin (Phalloidin, magenta) and (**F**) axons stained for tyrosinated (Tyr, red) and detyrosinated α-tubulin (dTyr, turquoise). Scale bar, 10 µm. Comparison of area **(G)** and tyrosinated α-tubulin signal **(H)** of the VASH1-deleted DRG growth cones treated or not with CNP. n/N: (**G**) 131/5 (Control) and 138/5 (CNP), (**H**) 100/3 (Control) and 99/3 (CNP) with n indicating number of growth cones and N number of E13 mice embryos. Data are expressed as mean ± SD and as fold change over control. Statistical significance was determined using Mann-Witney test. ns p > 0.05. a.u.: arbitrary unit

Taken together, these data suggest that CNP induces growth cone enlargement through VASH1, which thereby acts as a downstream mediator and target of PRKG1α.

### PRKG1α phosphorylates human VASH1 in vitro at Ser337 and impairs its detyrosination activity

To test whether PRKG1α phosphorylates VASH1, we resorted to an in vitro kinase assay based on protein thio-phosphorylation (Banko et al., 2011). In addition to the full-length recombinant VASH1-SVBP complex, we used truncated versions of the enzyme complex lacking either the N-terminal (CD+Ct) or C-terminal domains (Nt+CD), or all flanking regions of the core domain (CD) of VASH1 (previously used in (Ramirez-Rios et al., 2023), Figure S4A). In the presence of PRKG1α and the ATP-γ-S phosphate donor we could observe clear VASH1 thio-phosphorylation (Figure 4A). In these experimental conditions, the entire phosphorylation of VASH1 came from its C-terminal, as neither VASH1_CD nor VASH1_Nt+CD exhibited detectable thio-phosphorylation signals. These in vitro data showed that VASH1-SVBP complexes can be phosphorylated by PRKG1α and that the site(s) of phosphorylation are located on its C-terminal domain which we previously demonstrated is crucial for microtubule interaction (Ramirez-Rios et al., 2023).

**Figure 4.**
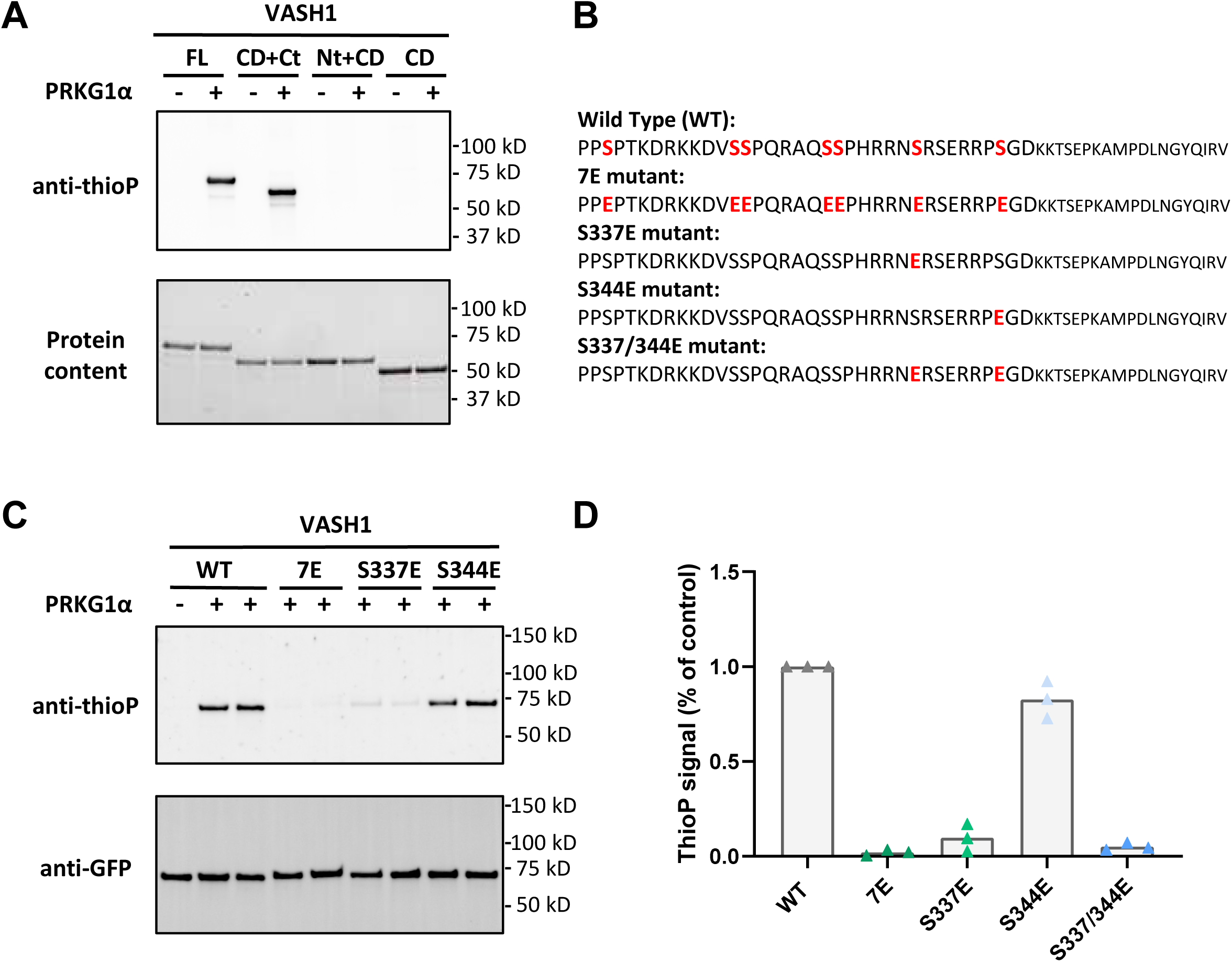
Recombinant VASH1 is phosphorylated by PRKG1α at its C-terminus on the serine 337. (**A**) Recombinant sfGFP-VASH1-SVBP (FL, full length) and truncated versions (CD, core domain; Nt, N-terminal region; Ct, C-terminal region, see Figure S4A) were incubated without or with PRKG1α (10 nM) and ATP-ϒS for 1 h and detected by immunoblot using an anti-thiophosphate ester antibody (anti-thioP). Protein content confirms similar loading of the samples. (**B**) C-termini of VASH1 and mutants developed in this study with the mouse- and human-conserved serine residues mutated or not to glutamates in red. (**C**) sfGFP-VASH1-SVBP (wild type enzyme, WT) along with different phosphomimics (S337E, S344E, S337/344E, and 7E, see **B**) were incubated without or with PRKG1α (10 nM) and ATP-ϒS for 1 h. Anti-GFP signal confirms equal loading of the samples. **(D)** The thio-phosphorylation signals of the different phosphomimic mutants are expressed relative to the WT enzyme of the same experiment. Three independent experiments were analyzed.

Our next objective was to pinpoint on the residues crucial for PRKG1α phosphorylation at this VASH1 C-terminal. This domain contains seven serine residues, which are conserved in humans and mice (Figure 4B, serines are in red). We carried out mass spectrometry (MS)-based proteomic analysis on samples of recombinant full-length VASH1-SVBP (V1 construct, Figure S4A) phosphorylated by PRKG1α in the presence of ATP (same conditions as set-up in the previous experiment but replacing ATP-γ-S with ATP). MS analyses were performed after Lys-C or trypsin digestion of the protein. Due to the highly basic nature of VASH1 C-terminus containing numerous lysines and arginines, peptides from this region are barely detectable (as previously reported (Aillaud et al., 2017)). Out of the seven conserved serines at the C-terminal of VASH1, triplicate analyses of the digests allowed to cover five residues, with no phosphorylation detected on any of them; S344 and S337 were not detected (Figure S4B, Table S1). Phospho-peptides enrichment did not improve coverage of phosphorylated serines in this domain of the protein (data not shown). Interestingly, the two serine residues not detected by MS analyses, S337 and S344, are predicted to lie within PRKG1 consensus motif XRRXSXX (Wood et al., 1996).

To obtain further insights, we created mutants to assess susceptibility to PRKG1α phosphorylation in vitro and to investigate how mimicking phosphorylation affects VASH1 activity. We previously generated the 7E recombinant mutant by replacing the seven C-terminal serine residues conserved in humans and mice with glutamate and showed that it cannot be phosphorylated by PRKG1α (Meyer-Jens M, 2026). Figure 4C reproduced our previous findings and narrowed our focus to the two best candidates by creating two single point-mutants, S337E and S344E, as well as the double mutant S337/344E (Figure 4B-D). Fluorescence immunoblots showed strong thiophosphate labelling for native VASH1, whereas the 7E mutant showed no signal. The S337E mutant showed a marked (∼90%) reduction in signal, while the S344E mutant exhibited only a modest (∼18%) decrease. The double mutant S337/344E showed near-complete loss (∼95%) of phosphorylation signal. Altogether these data implicate Ser337 as the dominant PRKG1α target site.

To determine whether phosphorylation at serine 337 affects VASH1 catalytic function, we compared the detyrosination activity of the wild-type (WT) enzyme and the mutant S337E, mimicking constitutively phosphorylated form, by immunofluorescence on taxol-stabilized microtubules (Figure 5). In the same experimental conditions, recombinant VASH1-SVBP enzyme showed a robust detyrosination of the microtubules characterized by a significant, widespread decrease in Tyr tubulin signal and a corresponding increase in dTyr signal. In contrast, the S337E mutant induced much lower changes in the α-tubulin types, consistent with reduced enzymatic activity (Figure 5A). Quantitative analysis of the activity of the recombinant proteins confirmed a significant difference in detyrosinated and tyrosinated tubulin signals for S337E mutants compared to wild type enzymes (Figure 5B), showing that phosphorylation at serine 337 inhibits the enzymatic activity of the VASH1-SVBP complex. The S337/344E double mutant showed similar reduction in activity to that of the single S337E mutant (Figure 5B), reinforcing serine 337 as the functionally dominant site. Together our data reveal that a single negative charge at the VASH1 C-terminus is deleterious to its catalytic activity.

**Figure 5.**
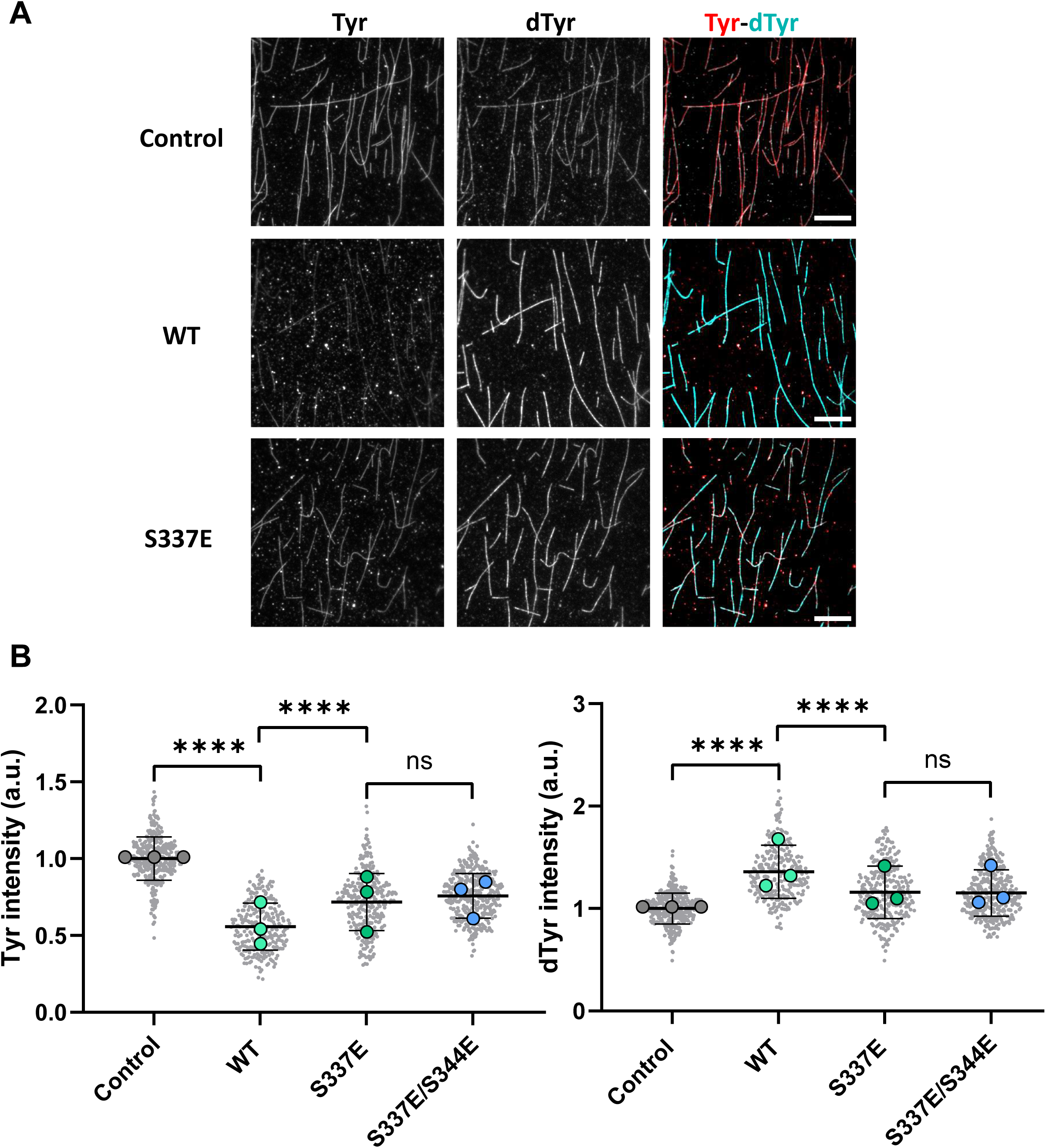
The S337E phosphomimetic of VASH1 has reduced detyrosination activity in vitro. Comparison of the activity of sfGFP-VASH1-SVBP (WT) and the mutant (S337E) enzyme complexes (50 pM) on taxol-stabilized HeLa tyrosinated (Tyr) microtubules measured by immunofluorescence in BRB40 buffer supplemented with 50 mM KCl as in (Ramirez-Rios et al., 2023). (**A**) Representative images of tyrosinated (Tyr, red) and detyrosinated (dTyr, turquoise) α-tubulin pools of microtubules after 30 min incubation in the absence (control) or presence of the indicated enzyme complexes. Scale bar, 10 μm. (**B**) Quantification of tyrosinated (Tyr) and detyrosinated (dTyr) tubulin signals. Each point represents a microtubule, at least 270 microtubules across 3 experiments done with 3 independent sets of recombinant enzymes were analyzed. Data are represented as mean ± SD. Statistical significance was determined using Kruskal-Wallis test, ****P < 0.0001.

To further investigate the mechanism underlying this functional impairment, we compared the microtubule-binding dynamics of VASH1-WT and VASH1-S337E using single-molecule total internal reflection fluorescence (TIRF) microscopy (Figure S5). Binding of GFP-tagged enzymes were visualized on tyrosinated microtubules over the first 10-15 min after enzyme addition, a time window chosen to minimize substrate depletion and allow accurate comparison of binding behaviour (Ramirez-Rios et al., 2023). Kymograph analysis revealed striking differences: VASH1-WT exhibited frequent, short-lived binding events along the microtubule lattice, whereas VASH1-S337E displayed sparse and punctate binding events (Figure S5A). Quantitative analysis of recombinant proteins showed that S337E approximately had a twofold reduction in residence time (0.14 s for S337E vs. 0.33 s for WT) and a 32% lower binding frequency compared to WT (Figure S5B, S5C), indicating a substantial decrease in microtubule engagement for the mutant.

Together, these findings identify serine 337 as a critical regulatory site whose phosphorylation by PRKG1α attenuates both the microtubule-binding affinity and the detyrosination activity of VASH1.

### The phosphorylated VASH1 form is detected in mouse DRG neurons and regulated through a CNP-PRKG1 axis

To further investigate the regulatory mechanism of VASH1 in cells, we developed a polyclonal antibody recognizing VASH1 phosphorylated at serine 337 in the human protein or at serine 347 in the mouse protein which contains 10 supplementary residues in its N-terminal domain (anti-P[S337]VASH1, see Material and Methods). The novel antibody was shown to be highly specific to the phosphorylated form of the enzyme using the in vitro phosphorylation assay (Figure S6A).

We first tested this antibody in immunofluorescence by comparing anti-P[S337]VASH1 signals from WT and *Vash1* KO neurons cultured from DRG explants (Figure 6A, B, S6B). A faint, sparse signal was observed in axons and growth cones of both WT and *Vash1* KO neurons. However, the signal intensity was much lower in *Vash1* KO than in WT neurons (∼33% and ∼22 % lower in growth cones and axons, respectively). This likely revealed the presence of the phosphorylated form of the enzyme on top of a substantial non-specific background signal in neuronal compartments. Treatment of WT neurons with CNP or DT-3 did not affect P[S337]VASH1 signal intensity in growth cones (Figure 6C) or along axons (Figure S6C), most likely due to the high background noise observed with this approach.

**Figure 6.**
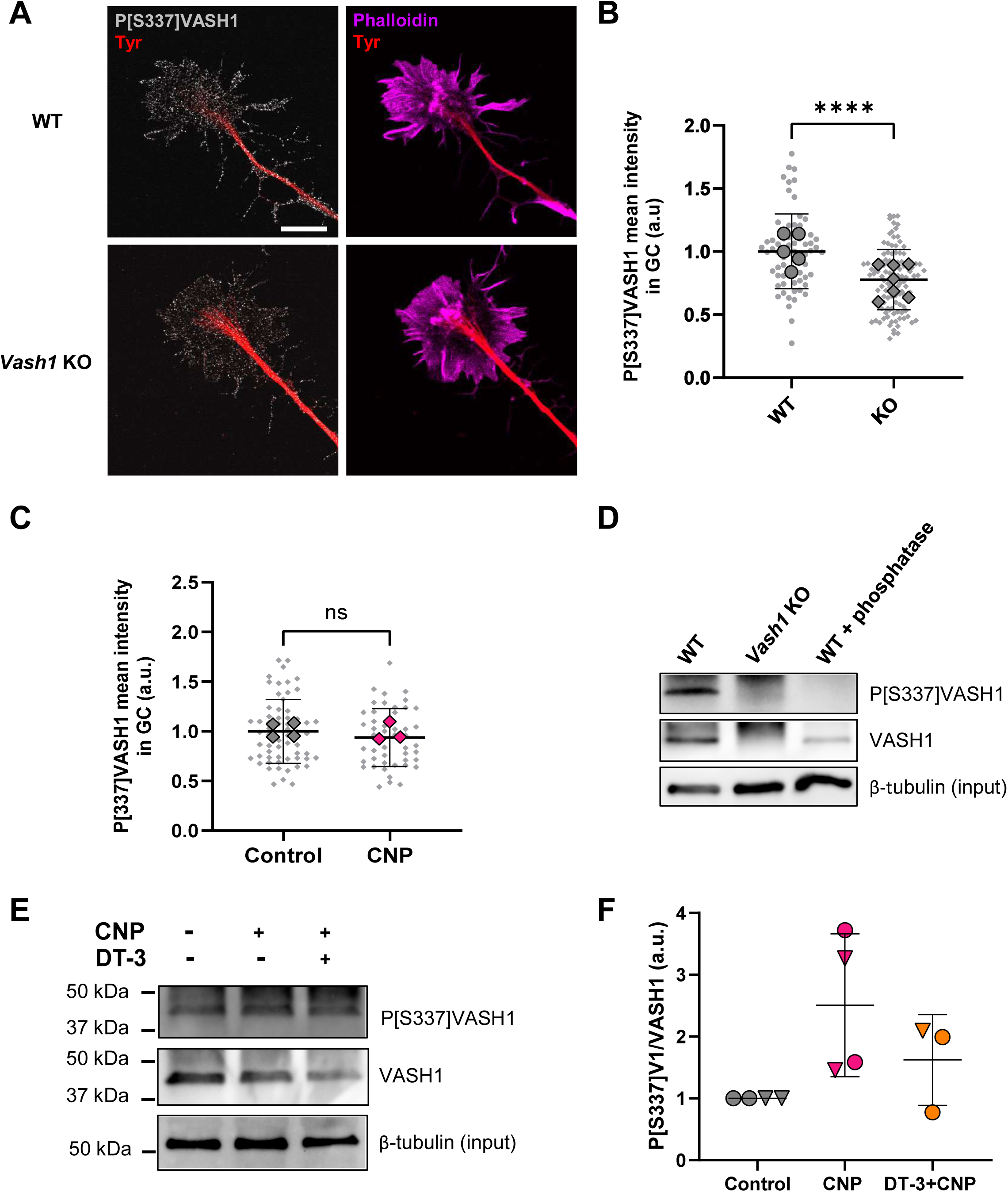
VASH1 is phosphorylated on C-terminal serine 337 under CNP treatment. **(A)** Representative confocal images of axon with growth cones stained for tyrosinated α-tubulin (Tyr, red), F-actin (phalloidin, magenta), and phosphorylated-VASH1 (P[S337]VASH1, gray). Scale bar, 10 µm. (**B**) Quantification of P[S337]VASH1 in growth cones of WT and *Vash1* KO neurons. n/N: 67/5 (WT), 111/7 (KO) with n indicating the number of growth cones and N of E13 embryos. Data are expressed as mean ± SD and as fold change over WT. (**C**) Quantification of P[S337]VASH1 in growth cones without treatment or with CNP treatment. n/N: 61/4 (control), 48/3 (CNP) with n indicating the number of growth cones and N the number of E13 embryos. Data are expressed as mean ± SD and as fold change over control. Statistical significance was determined using Mann-Witney test. **** p < 0.0001, ns p > 0.05. (**D, E**) Detection by immunoblot of VASH1 protein and its phosphorylated form in cultured DRG neurons after immunoprecipitation using an anti-SVBP antibody. β-tubulin staining shows similar protein content in the immunoprecipitation inputs. (**D**) Both VASH1 protein and its phosphorylated form are detected in WT but not in *Vash1* KO DRGs. Treatment with Lambda protein phosphatase results in the loss of P[S337]VASH1 band. (**E, F**) CNP induces phosphorylation of S337 of VASH1, which is reversed by DT-3, as shown by representative immunoblots from an experiment done with cultured DRG explants (**E**) and quantifications of the P[S337]VASH1/VASH1 ratios from 4 independent experiments (**F**). In **F**, triangles correspond to data from immunoprecipitation experiments done with cultured DRG dissociated neurons, and circle to immunoprecipitation experiments done with cultured DRG explants.

We then thought to analyse P[S337]VASH1 contents in DRG explants and dissociated neurons after enriching the detyrosinating enzyme by immunoprecipitation with the anti-SVBP antibody (Figure 1A) to achieve greater specificity. Figure 6D shows a band at the correct molecular weight with either the anti-VASH1 or anti-P[S337]VASH1 antibody in DRG neurons cultured from WT mouse but not from *Vash1* KO mouse. Additionally, enriched VASHs proteins from WT DRG neurons were treated with Lambda protein phosphatase, which led to the loss of the P[S337]VASH1 band confirming that the signal observed with the novel antibody corresponds to the expected phosphorylated form of VASH1. Next, cultured DRG explants or dissociated neurons were treated with CNP in the absence or presence of the PRKG1 inhibitor DT-3. After enrichment with anti-SVBP antibody, VASH1 and P[S337]VASH1 levels were analyzed by immunoblotting, and the P[S337]VASH1/VASH1 ratio was quantified (Figures 6E, F). Interestingly, CNP treatment consistently induced a significant increase in the proportion of phosphorylated VASH1, with the P[S337]VASH1/VASH1 ratio increasing between 1.5- and 3.7-fold. Moreover, the PRKG1α inhibitor DT-3 prevented the effect of CNP in both DRG explants and dissociated neuron preparations.

Altogether, these data based on use of a newly developed antibody, allowed to detect the presence of the S337-phosphorylated form of the VASH1 protein in DRG neurons and demonstrated that it accumulates following treatment with CNP in a PRKG1α-dependent mode.

## DISCUSSION

Early steps of DRG neuron development are regulated by C-type natriuretic peptide (CNP), which positively promotes axonal branching and growth through binding and activation of the NPR2 receptor, leading to increased cGMP levels and subsequent activation of PRKG1α (Schmidt et al., 2009; Zhao and Ma, 2009). While these upstream components of the regulatory signaling pathway have been characterized, the downstream mechanisms mediating these effects, particularly the targets phosphorylated by PRKG1α, have remained elusive. Evidence from previous work has highlighted a central role for microtubule dynamics in CNP-mediated regulation of DRG axon development, revealing that tubulin polymerization is required for the growth-promoting effects of CNP (Xia et al., 2013). This finding is consistent with the broader principle that, during neuronal development, signaling pathways triggered by guidance cues generally converge on the cytoskeleton to shape axonal morphology (Akiyama et al., 2016; Bouquet and Nothias, 2007; Dent and Gertler, 2003).

In the present study, we investigated the potential role of the CNP/cGMP/PRKG1 signaling cascade in regulating the detyrosination/re-tyrosination cycle of tubulin, a key determinant of microtubule dynamics, in DRG neurons. We show that CNP promotes growth cone enlargement, reduces detyrosinated tubulin along the axon and increases tyrosinated tubulin within the growth cone. These findings are consistent with previous observations that growth cone remodeling, considered a priming step for axon bifurcation in the spinal cord, is associated with increased levels of tyrosinated tubulin in cultured DRG neuron (Dumoulin et al., 2018a). We further revealed the presence of VASH1-SVBP detyrosination complex in DRG neurons, whereas VASH2-SVBP was undetected. Notably, pharmacological inhibition of vasohibins using epoY (Aillaud et al., 2017) produced effects comparable to those elicited by CNP, and CNP failed to exert its effects in VASH1-deficient DRG neurons. Taken together, these observations strongly suggest a model in which VASH1 is a likely substrate of PRKG1α in the downstream CNP signaling pathway.

We found that epoY had a slightly smaller effect than CNP on the size of the growth cone. Although we cannot exclude the possibility that this difference reflects incomplete inhibition of VASH activity by epoY, it may also indicate the involvement of additional downstream effectors of the CNP/cGMP/PRKG1 pathway. One potential candidate is GSK-3β, which can be phosphorylated by PRKG1. This phosphorylation inhibits the kinase, which disinhibits CRMP2 and promotes microtubule growth (Kim et al., 2006; Liz et al., 2014; Xia et al., 2013). In addition, a recent study indicated that the CNP/cGMP pathway could also control Ca^2+^ signaling and remodel the actin cytoskeleton in embryonic DRG neurons (Balmes et al., 2026).

Although no kinase regulating the activity of VASH1 had been identified to date, phosphorylation of VASH1 had been proposed as a potential mechanism that could modulate the interaction between VASH-SVBP enzymes and microtubules (Ramirez-Rios et al., 2023). The serine-rich VASH1 C-terminus, which is involved in microtubule binding, represents a plausible target for regulatory phosphorylation. Using in vitro assays, we demonstrated that PRKG1α can directly phosphorylate recombinant VASH1 enzyme on its C-terminal serine 337, thereby regulating its binding to microtubules and detyrosination activity. Notably, the introduction of a single negative charge at the VASH1 C-terminus site was sufficient to markedly alter VASH1 function, highlighting the sensitivity of this regulatory region. Finally, using a newly custom-made antibody, we demonstrated the presence of the corresponding phosphorylated form of VASH1, P[S337]-VASH1, in DRG neurons and revealed its modulation under CNP-treatment.

Collectively, our results support a model in which PRKG1α-mediated phosphorylation of VASH1 at serine 337 acts as a molecular switch that fine-tunes VASH1 activity, thereby modulating microtubule detyrosination and thus cytoskeleton dynamics within growth cone. The tubulin detyrosination/re-tyrosination cycle is linked to microtubule dynamics via the regulation of microtubule interactions with specific cellular effectors such as the depolymerising motor kinesin-13 (Sanyal et al., 2021). Whereas detyrosinated microtubules are resistant to this motor-driven depolymerization, tyrosinated microtubules are the preferred substrates (Peris et al., 2009). Within growth cone, dynamic tyrosinated microtubules are known to extend at the periphery along the actin filaments of filopodia enabling growth, environmental exploration and cues guidance (Baas and Black, 1990; Brown et al., 1993; Conde and Caceres, 2009; Kalil and Dent, 2005). Based on our findings, we propose that CNP-induced phosphorylation of VASH1 locally suppresses microtubule detyrosination, thereby increasing the abundance of tyrosinated tubulin and promoting microtubule extension into the growth cone. Such cytoskeletal remodeling could facilitate the morphological changes that precede DRG axon bifurcation. This is consistent with a previous study showing that tubulin re-tyrosination mediated by TTL enzyme is also essential for growth cone morphology and functionality in precerebellar nuclei neurons, with lack of TTL leading to abnormal cytoskeletal organization and impaired substrate adhesion of their growth cone (Marcos et al., 2009). TTL KO growth cones exhibit excessive enlargement with disoriented filopodia due to the infiltration of detyrosinated microtubules into the peripheral actin-rich domain. These observations support the notion that tyrosinated tubulin facilitates the crosstalk between microtubule entry and actin regulation, pivotal for maintaining growth cone morphology and facilitating axonal branching (Dent and Kalil, 2001).

Altogether, these studies show that enzymes governing both sides of the tubulin detyrosination/re-tyrosination cycle are implicated in the reorganization of the growth cone cytoskeleton. They underscore the critical role of a finely tuned and spatially regulated balance detyrosinated and tyrosinated tubulin in shaping neuron development. They also provide a further insight into the harmful neurodevelopmental phenotype observed in SVBP-deficient mice and human individuals, which lack the VASH detyrosinating enzymes (Pagnamenta et al., 2019). Patients with *SVBP* genetic variants exhibit significant delays in brain development and microcephaly, accompanied by symptoms such as mirror movements and spastic paraparesis, leading to impaired gait and motor function (Iqbal et al., 2019; Pagnamenta et al., 2019). It is plausible that at least part of these features, indicative of neuro-muscular defects, may arise from errors in peripheral neuron development.

This study reveals a novel CNP/cGMP/PRKG1α/VASH1 signalling cascade that directly links an extracellular developmental cue to the regulation of microtubule tyrosination and dynamics in DRG neurons (Figure 7). It was recently shown that neprilysin (an endopeptidase that inactivates several peptide hormones) inhibition reduces microtubule detyrosination in cardiomyocytes through the cGMP/PRKG1/VASH1 signaling axis (Meyer-Jens M, 2026). Together, these observations suggest that VASH1 may represent an essential and conserved downstream effector of natriuretic peptide signaling across different cell types. More broadly, these findings provide a molecular and mechanistic explanation on how natriuretic peptides signaling can regulate cytoskeletal organization, and extend our current understanding of kinase-mediated regulation of the cytoskeleton to the tubulin code. By identifying VASH1 phosphorylation as a mechanism linking extracellular cues to tubulin detyrosination, this work highlights a previously unrecognized signaling pathway that influences microtubule dynamics to shape neuronal development.

**Figure 7.**
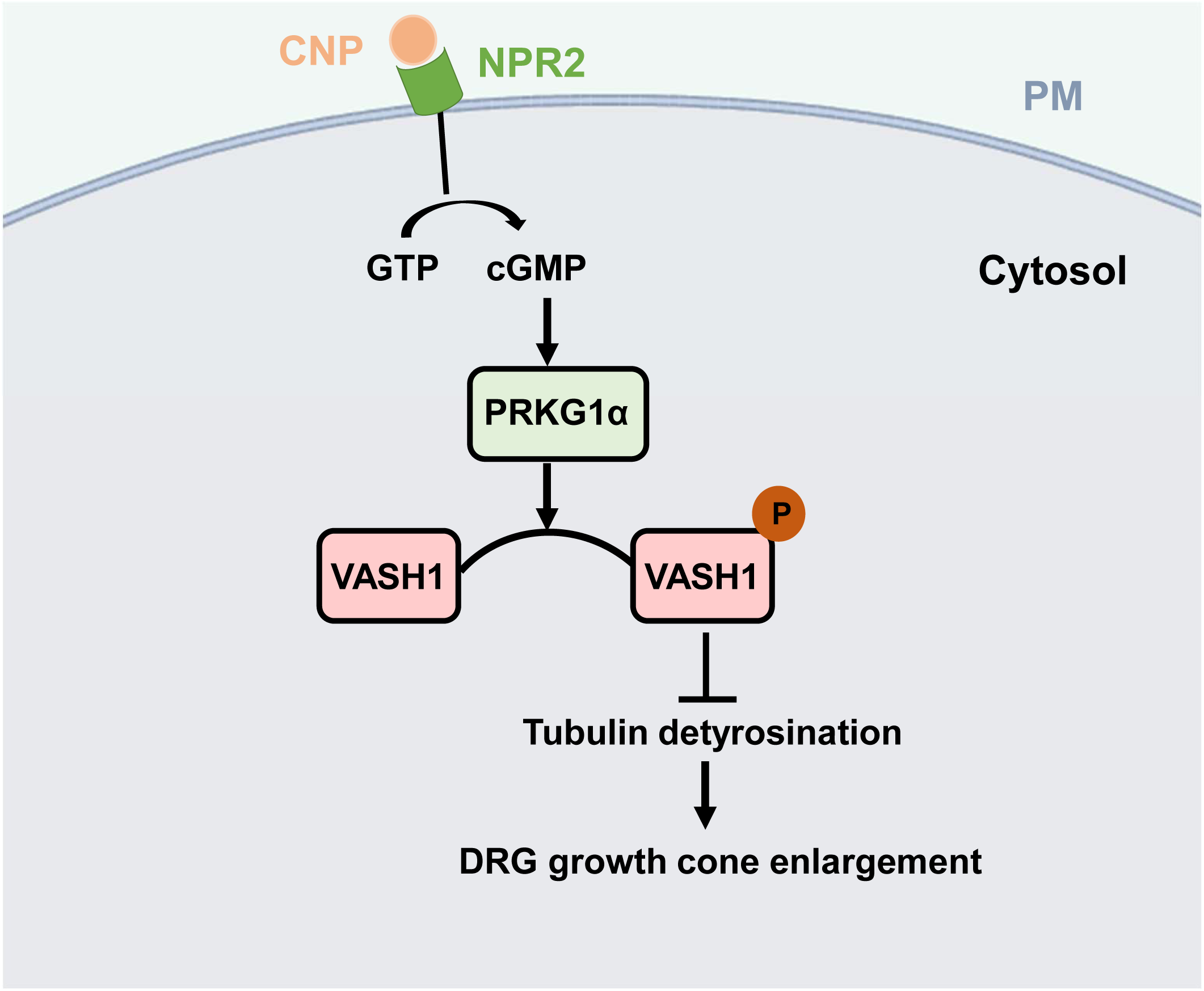
Schematic illustration of CNP/cGMP/PRKG1α pathway involved in VASH1 mediated regulation of microtubule detyrosination in DRGs development.

## MATERIALS AND METHODS

### Animals and gene editing

The study protocol complied with the guidelines of the French Ministry of Research and Directive 2010/63/EU of the European Parliament on the protection of animals used for scientific purposes. Mice were housed in cages with environmental enrichment, in a temperature-controlled room with a 12-hour light/dark cycle with ad libitum access to water and food. Every precaution was taken to minimize stress and the number of animals used in each experiment. C57/Bl6 mice (Janvier) were generally used as wild-type mice, except for immunoprecipitation experiments where CD1/Swiss mice (Janvier) were also employed.

The *Vash1* KO was generated by CRISPR/Cas9 editing. Guide selection and off-target predictions were made with CRISPOR software (Haeussler et al., 2016). Target sequences (CCACTCTGGCACTAAACGTT on the coding strand of *Vash1)* was unique in the genome, while predicted off-target sequences on the same chromosome were all non exonic and with at least 4 mismatches with on-target sequences, partly localized close to the PAM. A dual guide RNA was prepared by combining synthetic tracrRNA (TriLink BioTechnologies) and crRNA (Eurogentec, Angers, France). Sequences are given in Supplementary Table S2. A cloning free procedure was used to modify the *Vash1* locus, according to (Markossian et al., 2018), by microinjection of Cas9/RNA complexes into *in vitro* fertilized B6D2F1xFVB embryos. F0 mosaic animals born after reimplantation of microinjected embryos were genotyped by PCR/Sanger sequencing and then mated to C57BL/6J mice to provide F1 founder mice. These mice were selected according frameshifts introducing as less as possible extra amino acids. The allele was designated Vash1T38*, and the corresponding VASH1 mutant protein sequence was MPGGKKVVPSGSSSASPNAAATTTAAAAAAAAAPHSG. At least 6 additional back-crosses were performed with C57BL/6J mice before phenotyping in order to avoid possible confounding effects due to off-target mutations. Genotyping was performed by PCR amplifications on alkaline lysates from small ear lobe samples.

### Embryonic DRG explants and dissociated neurons cultures

For explants, coverslips (immunofluorescence) or plastic dishes (immunoprecipitation) were coated overnight with poly-L-lysine (0.5 mg/mL, Sigma), washed with sterile water and subsequently coated with laminin (20 µg/mL, Sigma) for 3-4 h at room temperature and maintained in Hibernate-E medium (Gibco). DRGs were dissected from wild-type (WT) and *Vash1* knockout (KO) mouse embryos at embryonic day 13 (E13) in Hibernate-E medium. They were then transferred on coated coverslips or dishes in a small volume of 0.4% methylcellulose, B-27 supplement, L-glutamine in hibernate medium. For immunofluorescence, three to four DRG explants were transferred onto each coated coverslip; for immunoprecipitation, ∼80 DRGs explants were transferred onto each coated dish. Then culture medium containing neurobasal medium (Gibco), B-27 supplement (Gibco), L-glutamine (Gibco), Penicillin-Streptomycin (Gibco), and nerve growth factor (10 ng/mL, Alomone Labs) was added.

For dissociated neurons, plastic dishes were coated overnight with poly-L-lysine (0.5 mg/mL, Sigma), washed with sterile water, and subsequently coated with laminin (3 µg/mL; Sigma) for 2 h at 37°C. Dishes were then air-dried under sterile conditions. DRGs were dissected in Hibernate medium, collected by centrifugation, resuspended in trypsin-EDTA (0.05%, Gibco) and incubated for 22 min at 37°C. Trypsinization was stopped by adding Dulbecco’s Modified Eagle Medium (DMEM) supplemented with 10% horse serum. After centrifugation, the cells pellet resuspended in a medium consisting of Neurobasal Plus medium (Gibco), B-27 supplement, GlutaMAX (Gibco), penicillin-streptomycin, and nerve growth factor (50 ng/mL) were mechanically dissociated by trituration and plated in dishes with additional culture medium. Dissociated cells from ∼100 DRGs were used per immunoprecipitation.

DRG explants or dissociated neurons were then maintained 20-22h at 37°C in a humidified atmosphere of 95% air and 5% CO₂ in the culture medium and then incubated without or with 500 nM CNP (Sigma) for 1 h following or not a pretreatment with a PRKG1α inhibitor (DT3 at 150 nM, Sigma) for 2 h or a VASHs inhibitor (epoY 50 µM, (Aillaud et al., 2017)) for 3 h.

### Immunofluorescence on DRG explants

DRG Explants were fixed in 4% paraformaldehyde and 1.5% sucrose in phosphate-buffered saline (PBS) for 20 min at room temperature, followed by PBS washes. Samples were permeabilized with 0.1% Triton X-100 (Sigma-Aldrich) in PBS for 4-5 min, then washed again with PBS. For growth cone area and tyrosinated (Tyr) α-tubulin intensity measurements, neurons were stained with rat anti-tyrosinated tubulin (YL1/2, 1:3000), followed by Cy3-conjugated anti-rat IgG (Jackson ImmunoResearch, 1:500) and F-actin was labeled with phalloidin–ATTO 647 (ATTO-TEC, 1:500). For tubulin intensity measurements (Tyr, dTyr) in axons, primary antibodies included rat anti-tyrosinated tubulin (YL1/2, 1:3000) and rabbit anti-detyrosinated tubulin (1:1000) (Aillaud et al., 2016). Secondary antibodies were Alexa Fluor 488-conjugated anti-rat IgG and Cy3-conjugated anti-rabbit IgG (Jackson ImmunoResearch, both at 1:500). For detection of phosphorylated species of VASH1 in DRGs, primary antibodies were rabbit P[S337]VASH1(1:1000) and rat anti-tyrosinated tubulin (YL1/2, 1:3000). Secondary antibodies were Alexa Fluor 488-conjugated anti-rabbit IgG and Cy3-conjugated anti-rat IgG (Jackson ImmunoResearch, both at 1:500). Primary antibodies were incubated for 2 h at room temperature, followed by three washes in PBS-0.1% Tween-20. Secondary antibody incubation was performed for 1 h at room temperature, followed by three additional washes in PBS-0.1% Tween-20. Explants were mounted in DAKO mounting medium (Sigma-Aldrich) supplemented with Hoechst 33258 (1 µg/mL, Invitrogen). Images were acquired using a Zeiss LSM 710/Airyscan confocal microscope equipped with a 63× (NA 1.4) objective, using or not the Airyscan function.

### Constructs and enzymes purification for TIRF and immunofluorescence studies

The constructs in pETDuET-1 vector allowing production of human full-length sfGFP-VASH1-SVBP (V1_FL) and its truncated versions V1_CD, V1_Nt+CD and V1_CD+Ct were described in (Ramirez-Rios et al., 2023). The constructs allowing production of its mutated versions was obtained by PCR amplification with overlapping primers encoding substitutions S313E, S323E, S324E, S330E, S331E, S337E, S344E (mutant 7E, (Meyer-Jens M, 2026)), or S337E, or S344E, or both S337E and S344E (S337/344E mutant) in the C-terminal region of VASH1. Primer sequences are given in Supplementary Table S1. PCR was performed with Phusion DNA polymerase and circularization by In-Fusion HD Cloning kit. Coding regions were verified by DNA sequencing (Genewiz).

Protein expression and purification of all the His-tagged VASH1-SVBP complexes were performed as previously described in (Ramirez-Rios et al., 2023). Concentration of soluble proteins was measured using Bradford assay with BSA as the standard. All proteins were kept in storage buffer (20 mM Tris-HCl at pH 7.4, 150 mM NaCl, 2 mM DTT) and stored in liquid nitrogen until use. Active versions of the enzymes were employed except in Figure 4A, where dead versions (catalytic cysteine mutated to alanine, (Aillaud et al., 2017)) were used.

### In vitro kinase assay

In vitro thiophosphorylation-based kinase assay on VASH1 enzymes was carried out using a method adapted from (Banko et al., 2011). sfGFP-VASH1-SVBP (2 µM) proteins were incubated for 1 h at 30°C in the presence or absence of PRKG1α (10 nM, Sigma Aldrich Merck) in a buffer containing 40 mM Tris-HCl pH 7.5, 100 mM NaCl, 10 mM MgCl_2_, 5% glycerol, 1 mM dithiothreitol and 60 μM ATPγS (Abcam). The kinase reaction was stopped by addition of 20 mM EDTA. Alkylation was then performed by adding 50 mM PNBM (p-nitrobenzyl mesylate, Abcam) for 1 h and stopped with the addition of Laemmli buffer. Phosphorylation of proteins was detected in immunoblot analysis using an anti-thiophosphate ester antibody (Abcam).

To test the novel anti-P[S337]VASH1 antibody, in vitro phosphorylation of VASH1 was performed in the same conditions as above but in the presence of ATP instead of ATPγS. Phosphorylated VASH1 protein was then detected by immunoblot (see below).

### Preparation of tubulin and Taxol-stabilized microtubules

Brain tubulin purification and its labeling with either biotin or ATTO-565 fluorophore (ATTO-TEC Gmbh) were done as in (Ramirez-Rios et al., 2017). Tubulin from HeLa cells were produced and purified as previously described (Souphron et al., 2019) with some modifications. HeLa S3 cells (CLS Cell Lines Service GmbH) were first cultivated ∼5 days in adherent cultures, and then used to inoculate 2 liters of suspension culture that were cultured during ∼14 days at 65 rpm, 37°C, 5% CO_2_ in a New Brunswick S41i incubator (Eppendorf). Cells were then harvested, resuspended in a BRB80 buffer containing 80 mM PIPES (piperazine-N, N′-bis (2-ethanesulfonic acid))-KOH at pH 6.8, 1 mM MgCl_2_, 1 mM EGTA supplemented with 1 mM 2-mercaptoethanol, 1 mM PMSF (Phenylmethanesulfonyl fluoride), cOmplete EDTA-free protease inhibitor cocktail tablets (GE Healthcare) and 0.2 % (vol/vol) Triton X-100, and lysed by sonication. The cells lysate was clarified by centrifugation at 150 000 g. The supernatant obtained was then subjected to two rounds of polymerisation/depolymerization cycles (one in a low molarity buffer containing 0.08 M PIPES, the other in high-molarity buffer containing 0.5 M PIPES, both with GTP and glycerol) as described in (Souphron et al., 2019). The polymerised microtubules were finally spined down at 150 000 g for 30 min at 30° C. The pellet obtained was depolymerised on ice using cold BRB80 medium (20-30 min), followed by a centrifugation at 200 000 g for 20 min at 4° C. The supernatant obtained after this step contains exclusively tyrosinated tubulin. Taxol-stabilised microtubules were prepared as described in (Ramirez-Rios et al., 2023). Microtubules were polymerized using 45 μM tubulin (composed of 65% of HeLa tyrosinated tubulin, 30% biotinylated brain tubulin, and 5% ATTO-565–labeled brain tubulin) in BRB80 supplemented with 1 mM GTP. After microtubule formation, 100 μM Taxol were added and incubation was continued for 30 min. Tyr-microtubules were then centrifuged 10 min at 200,000 g and resuspended in BRB80 supplemented with 10 μM Taxol.

### TIRF in vitro assay

TIRF imaging assays were performed as previously described in (Ramirez-Rios et al., 2023). Taxol-stabilized microtubules enriched in Tyr tubulin were perfused into TIRF chambers and incubated at room temperature for 5 min. Unbounded microtubules were removed by three washes of 100 µl of BRB80 supplemented with 1% bovine serum albumin (BSA) and 10 μM Taxol. Finally, 30 µl of a solution containing 50 pM of the sfGFPVASH1-SVBP complex in BRB80 supplemented with 82 µg/ml catalase, 580 µg/ml glucose oxidase, 1 mg/ml glucose, 4 mM DTT, 0.5 mM Taxol, 0.01% methylcellulose (1,500 cp) were perfused. The chamber was sealed, and images were recorded within the first 30 min following addition of the assay-mix solution on an inverted microscope (Eclipse Ti, Nikon). The microscope was equipped with a Perfect Focused System, a CFI Apochromat TIRF 100×/1.49 N.A. oil immersion objective (Nikon), a warm stage controller (Linkam Scientific) and a Technicoplast chamber to maintain the temperature, an objective heater (OkoLab), an iLas2 TIRF system (Roper Scientific), and a sCMOS camera (Prime95B, Photometrics) controlled by MetaMorph software (version 7.10.3, Molecular Devices). For dual view imaging, an OptoSplit II bypass system (Cairn Research) was used as image splitter, and illumination was provided by 488- and 561-nm lasers (150 and 50 mW, respectively). Temperature was maintained at 35°C for all imaging purposes. Acquisition rate was one frame each 50 ms exposure (in streaming acquisition) during 45 s. TIRF movies were taken during the first 15 minutes following addition of enzyme complexes to microtubules (approximately 4-5 films could be taken within this time frame).

### In vitro detyrosination activity assay by immunofluorescence

VASH1 activity was measured as previously described (Ramirez-Rios et al., 2023). Taxol-stabilized microtubules were perfused in the same perfusion chambers and TIRF experiments conditions as above. After the addition of VASH1-SVBP enzymes (50 pM), chambers were incubated at 37°C for 30 min followed by three washes with 10 µM Taxol and 1% BSA in BRB80 (wash buffer). The incubation with primary antibodies (rat anti-tyrosinated tubulin (YL1/2, anti-Tyr, 1:6.000) and rabbit anti-detyrosinated tubulin (anti-dTyr, 1:1,000) was done for 15 min, followed by three washes with 100 µl of wash buffer. Secondary antibodies (anti-rat coupled to Alexa Fluor 488 and anti-rabbit coupled to Cyanine 3 from Jackson ImmunoResearch, both diluted to 1:500) were then added and incubated for 15 min, followed by three washes with 100 µl of wash buffer. Images were obtained immediately after using a LEICA DMI600/ROPER microscope controlled by Metamorph Video software.

### Quantification and Data Analysis

Growth cone morphology and fluorescence intensity were quantified using phalloidin staining to generate a binary mask. A custom ImageJ macro was used to measure the surface area and intensities of the different stainings within the growth cone.

Because growth cone size varied across experiments, areas were normalized to the mean growth cone area of the control condition within each experiment. Statistical analyses were performed using GraphPad Prism 8 (see Legends). Data were analyzed as indicated in Legend to Figures. Significance levels are indicated as follows: ns, not significant (p > 0.05), *p<0.5, **p < 0.01, ***p<0.001, ****p < 0.0001.

Tyrosinated (Tyr) and detyrosinated (dTyr) tubulin levels were quantified by manual tracing of 60 µm long lines over the axon starting at the growth cone front, and intensity values for Tyr- and dTyr-tubulin were measured. For each experimental condition, neurons from two replicate slides were analyzed. Mean intensity values were normalized to the mean intensity of the control group within each experiment.

Immunofluorescence and tracking of single VASH1-SVBP molecules for estimation of activity and binding parameters were analyzed using FIJI software and a homemade plugin, respectively, as in (Ramirez-Rios et al., 2023).

### Immunoprecipitation

Home-made anti-SVBP (anti-SVBP Gre, described in (Ramirez-Rios et al., 2023) was coupled to protein-G magnetic beads (Cytiva) using bis(sulfosuccinimidyle)suberate (ThermoFisher Scientific) as a crosslinker. DRG explants (∼80/immunoprecipitation) or neurons (dissociated from ∼100 explants/immunoprecipitation) isolated from E13 mice were lysed in 300 µL of buffer containing 80 mM PIPES-KOH (pH=6.7), 1 mM MgCl_2_, 1 mM EGTA, 150 mM NaCl, 1% NP-40, supplemented with cOmplete, EDTA-free Protease Inhibitor tablets (Roche) and PhosSTOP tablets (Roche; except in the experiment using phosphatase). The lysate was then incubated with the anti-SVBP-coupled protein-G magnetic beads, and the beads were rinsed 3-times with the lysis buffer. For phosphatase treatment experiments, bead-bound complexes were incubated with phosphatase (Lambda protein phosphatase, New England Biolabs) for 30 min at 30°C according to the manufacturer’s protocol before elution. The VASH-SVBP complexes were eluted with 100 mM glycine at pH=2.7, and the eluate was immediately neutralized with 1 M Tris (pH 8.0). Laemmli buffer was then added for subsequent protein analysis.

### Production of polyclonal anti-P[S337]VASH1

A novel antibody for human and mouse VASH1 phosphorylated at serine 337 or S347, respectively, was produced in rabbit by using the peptide HRRN**pS**RSERRP+C (p, phosphoryl group), acetylated at the N-terminal and linked at the N-terminal to the keyhole limpet hemocyanin protein via a cysteine. This sequence corresponds to amino acids 333-343 of human VASH1 and amino acids 343-353 of mouse VASH1. Rabbit serum was purified by differential cleaning using the non-phosphorylated and the phosphorylated peptides. Validation of these antibodies was performed by the in vitro kinase assay as described above using recombinant VASH1-SVBP complex in the presence or absence of the PRKG1 enzyme (Sigma) and ATP instead of ATPγS (Figure S6A). By convention, this antibody is named after the human form, P[S337].

### SDS-PAGE and immunoblotting

Recombinant proteins were separated on 4–20% gradient polyacrylamide gels (Bio-Rad) and transferred onto nitrocellulose membranes using the Trans-Blot Turbo Transfer System (Bio-Rad). Proteins immunoprecipitated from DRGs were separated on 4–15% gradient acrylamide gels (Bio-Rad) and transferred onto PVDF membranes. Membranes were blocked for 1 h in PBS containing 5% milk and 0.1% Tween-20. Primary antibodies include rabbit anti-thiophosphate ester (Abcam, 1:5000) and chicken anti-GFP antibodies (Millipore, 1:5000) for the in vitro kinase assays, and mouse anti β-tubulin (β3A11, (Aillaud et al., 2016); 1:10000), rabbit anti-VASH1 (VASH1 Gre, 1:10000, described in (Chen et al., 2020), rabbit anti-VASH2 (VASH2 Gre, 1:2000; described in (Ramirez-Rios et al., 2023)), and rabbit P[S337]VASH1 (1:2000, see above) for immunoprecipitations, and rabbit anti-GFP antibody (Jackson, 1:10000) and anti**-**P[S337]VASH1 (see above, 1:1000) to test anti**-**P[S337]VASH1 specificity. After washing, membranes were incubated with HRP-conjugated and/or fluorescence secondary antibodies (anti-rabbit and anti-mouse HRP antibodies, Jackson ImmunoResearch; anti-Chicken Alexa Fluor 488, Invitrogen; anti-rabbit Cyanine 5, Molecular Probes) and revealed using ECL Western blotting substrate or enhanced chemiluminescence substrates (Thermo Scientific). Signals were detected using a ChemiDoc™ MP Imaging System (Bio-Rad). Band intensities were quantified using Image Lab software (Bio-Rad), and statistical analyses were performed using GraphPad Prism.

### Mass spectrometry-based proteomic analyses

VASH1 phosphorylation was performed in the same conditions as for the in vitro kinase assays but in the presence of ATP instead of ATPγS, proteins separated on SDS-PAGE. The proteins in the cut gel bands corresponding to VASH1 (n=3) were digested in-gel using trypsin (modified, sequencing purity, Promega) or Lys-C (Mass Spec Grade, Promega), as previously described (Casabona et al., 2013), except that Tris(2-carboxyethyl)phosphine hydrochloride was used instead of dithiothreitol. The resulting peptides were analyzed by online nanoliquid chromatography coupled to MS/MS (Ultimate 3000 RSLCnano and Q-Exactive HF or Q-Exactive Plus, Thermo Fisher Scientific, for respectively two first and third replicates). For this purpose, the peptides were sampled on a precolumn (300 μm x 5 mm PepMap C18, Thermo Scientific) and separated in a 75 μm x 250 mm C18 column (Aurora Generation 2, 1.6 μm, IonOpticks) using a 20 min acetonitrile gradient. The MS and MS/MS data were acquired with Xcalibur (Thermo Fisher Scientific).

Peptides and proteins were identified by Mascot (version 3.1.0, Matrix Science) through concomitant searches against the sequence of GFP-Vash1-myc-His, the Uniprot database (*Escherichia coli* K12 taxonomy, 202402 version), and a homemade database containing the sequences of classical contaminant proteins found in proteomic analyses (keratins, trypsin, etc.). No enzyme specificity was set. Precursor and fragment mass error tolerances were set at respectively at 10 and 20 ppm. Peptide modifications allowed during the search were: Carbamidomethyl (C, fixed), Acetyl (Protein N-term, variable), Oxidation (M, variable) and Phosphorylation (S/T/Y, variable). The Proline software ((Bouyssie et al., 2020), version 2.2) was used for the compilation, grouping and filtering of the results (conservation of rank 1 peptides, peptide length ≥ 6 amino acids, false discovery rate of peptide-spectrum-match identifications < 1% (Coute et al., 2020), and minimum of one specific peptide per identified protein group). Proline was then used to perform a spectral counts-based analysis of the identified protein groups in the different samples.

The complete mass spectrometry proteomics data have been deposited to the ProteomeXchange Consortium via the PRIDE (Perez-Riverol et al., 2022) partner repository with the dataset identifier *PXD0XXXXX*.

## ACKNOWLEDGEMENTS

We thank all the zootechnicians of the Grenoble Institut des Neurosciences for animal care. We thank Homaira Nawabi and Sylvie Gory for help in setting-up in our team the DRG explants and dissociated neurons cultures, respectively. We acknowledge the contribution of SFR Biosciences (UMS3444/CNRS, US8/INSERM, ENS de Lyon, UCBL) facility: AniRA-PBES (Plateau de Biologie Expérimentale de la Souris, ENS de Lyon), especially S. Markossian, M. Teixeira, F. Henry and JL. Thomas.

This work was supported by the Leducq Foundation, research grant n° 20CVD01 (to M.-J. Moutin and L. Carrier), by Agence National de la Recherche (ANR), grant SPEED-Y n° ANR-20-CE16-0021 (to M.-J. Moutin). This work used the Photonic Imaging Center of Grenoble Institute of Neuroscience (PIC-GIN, University Grenoble Alpes – INSERM U1216), which is part of the ISdV core facility and certified by the IBiSA label. The proteomic experiments were partially supported by Agence Nationale de la Recherche under projects ProFI (Proteomics French Infrastructure, ANR-10-INBS-08 & ANR-24-INBS-0015) and GRAL, a program from the Chemistry Biology Health (CBH) Graduate School of University Grenoble Alpes (ANR-17-EURE-0003). Part of the mice were bred at the animal facility of IRIG-DRF-CEA, which is supported by funding from GRAL, a program of the University Grenoble Alpes graduate school, CBH-EUR-GS (ANR-17-EURE-0003).

---

**Figure S2.**
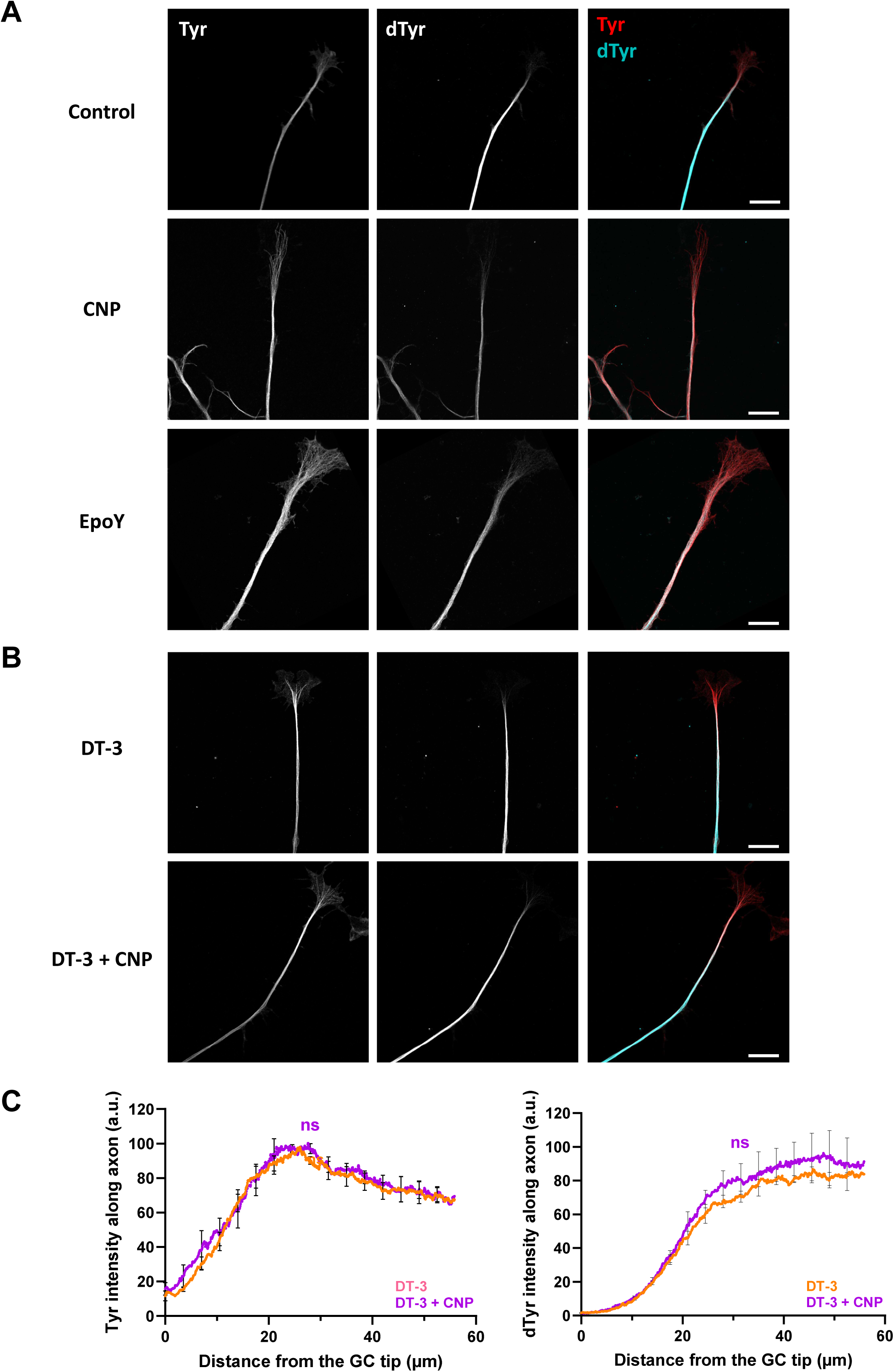
Effect of CNP, epoY and DT-3 on tyrosinated/detyrosinated α-tubulin levels in the axon of DRG neurons. (A, **B)** Representative confocal Airyscan images of axons stained for tyrosinated (Tyr, red) and detyrosinated α-tubulin (dTyr, turquoise) in the indicated conditions, with (A) same conditions as in Figure 2B and (B) same conditions as in Figure 1F. Scale bar, 10 µm. **(C)** Quantification of tyrosinated (Tyr) and detyrosinated (dTyr) α-tubulin along the axons. n/N: 95/6 (DT-3), 85/6 (DT-3+CNP) with n indicating number of growth cones and N number of slides (2 per experiment). Data are expressed as mean ± SEM (using N) and relative to data in the absence of CNP. Statistical significance was determined using two-way ANOVA and Sidak’s multiple comparison test. ns p > 0.05.

**Figure S3.**
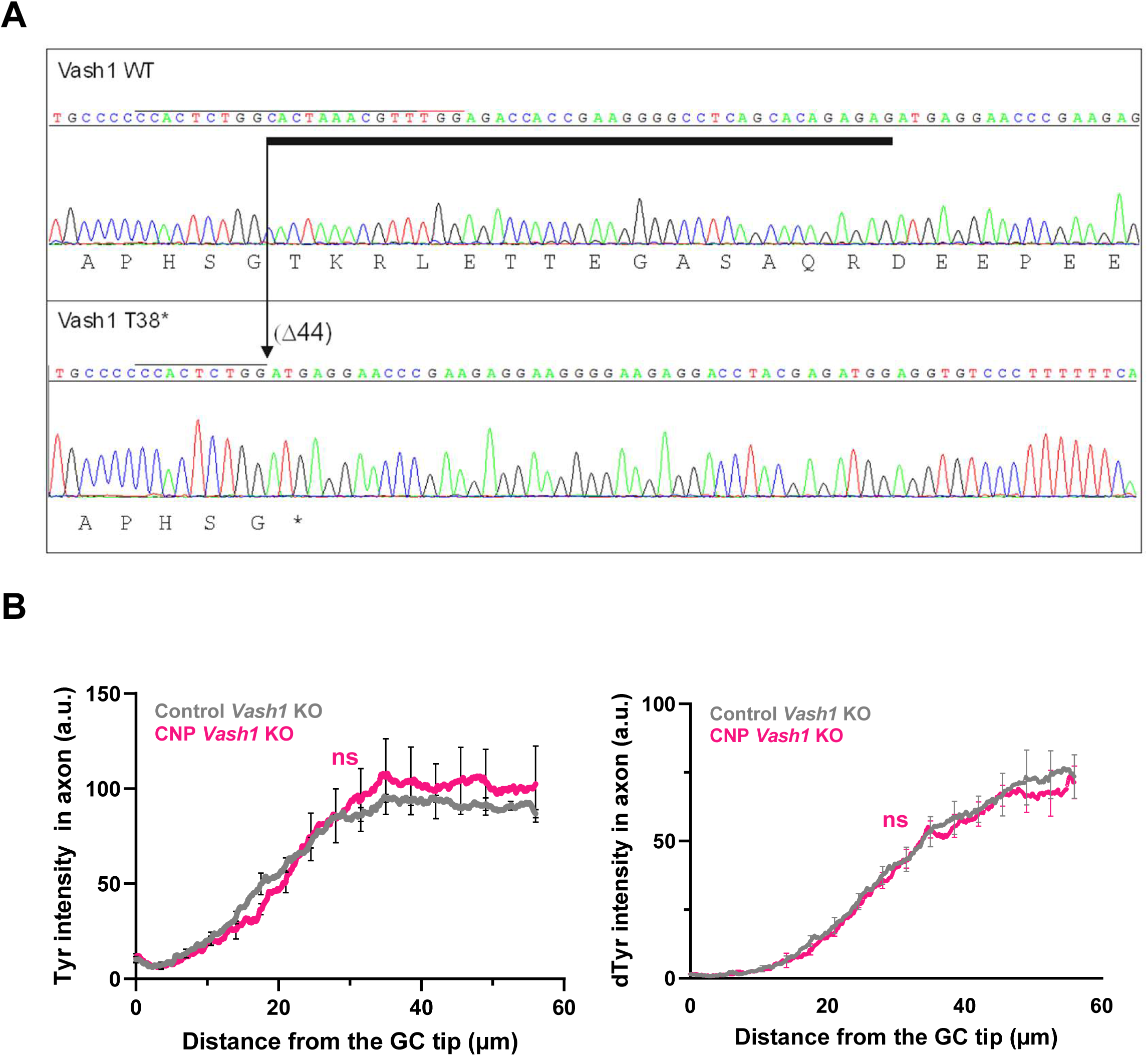
Vash1 knockout genotyping and loss of Tyr/dTyr axonal changes induced by CNP in the Vash1 KO. **(A)** *Vash1* gene DNA Sanger sequencing of control C57BL/6 mouse (WT) and of knock-out mouse Vash1 (T38*). The frameshift immediately introduces a stop codon. Black and red lines above the nucleotide sequence respectively indicate the positions of the crRNA and the protospacer adjacent motif (NGG), located on the coding strand of *Vash1*. Deletion site (arrow) and extent of the deletion are indicated above the WT sequences by a black bar. Translation product of Vash1 is shown below the electrophoregrams. (**B**) Quantification of tyrosinated (Tyr) and detyrosinated (dTyr) α-tubulin levels along the axons of DRG neurons from V*ash1* KO embryos in the absence or presence of CNP treatment. n/N: 76/6 (Control), 75/6 (CNP) with n indicating number of growth cones and N number of slides (2 per experiment). Data are expressed as mean ± SEM (using N) and relative to the wild type. Statistical significance was determined using two-way ANOVA and Sidak’s multiple comparison test. ns p > 0.05.

**Figure S4.**
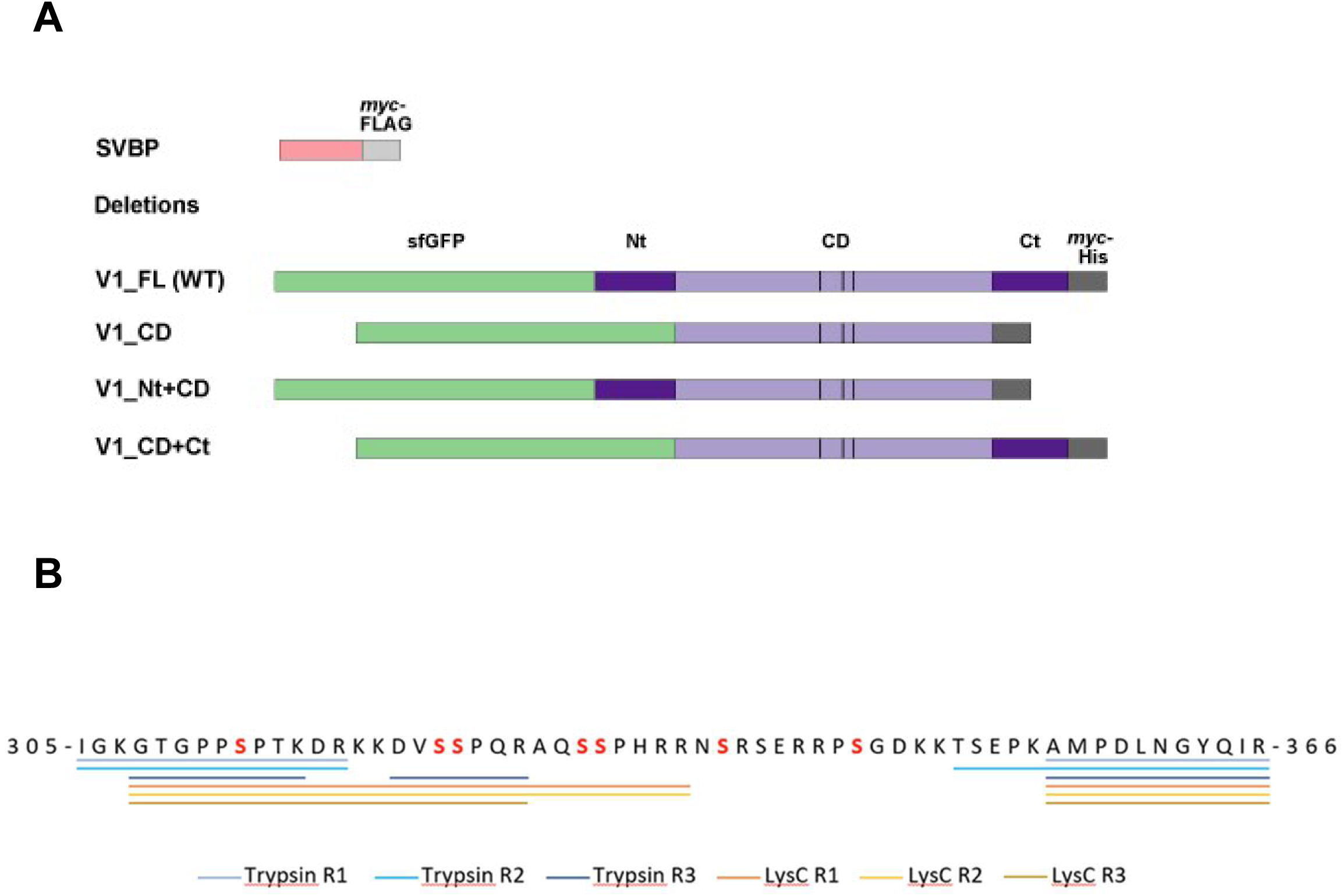
Recombinant proteins used in the study and sequence coverages of the C-terminal of recombinant VASH1 subjected to PRKG1α after analysis. (**A**) Schematic representation of the recombinant proteins used in this study with VASH1 (V1) in blue, SVBP in purple, sfGFP in green, tags (myc, FLAG, His) in gray. Vertical bars, catalytic triads. FL, full length; CD, core domain; Nt, N-terminal region; Ct, C-terminal region. (**B**) Sequence of the C-terminus of VASH1 with all the peptides identified in three independent purifications of the recombinant protein digested with either Trypsin or LysC. Peptides identified are represented with different coloured underlines. Serine which can potentially be phosphorylated by PRKG1α residues are indicated in red in the VASH1 sequence. No peptide carrying serine residues 337 and 344 were identified.

**Figure S5.**
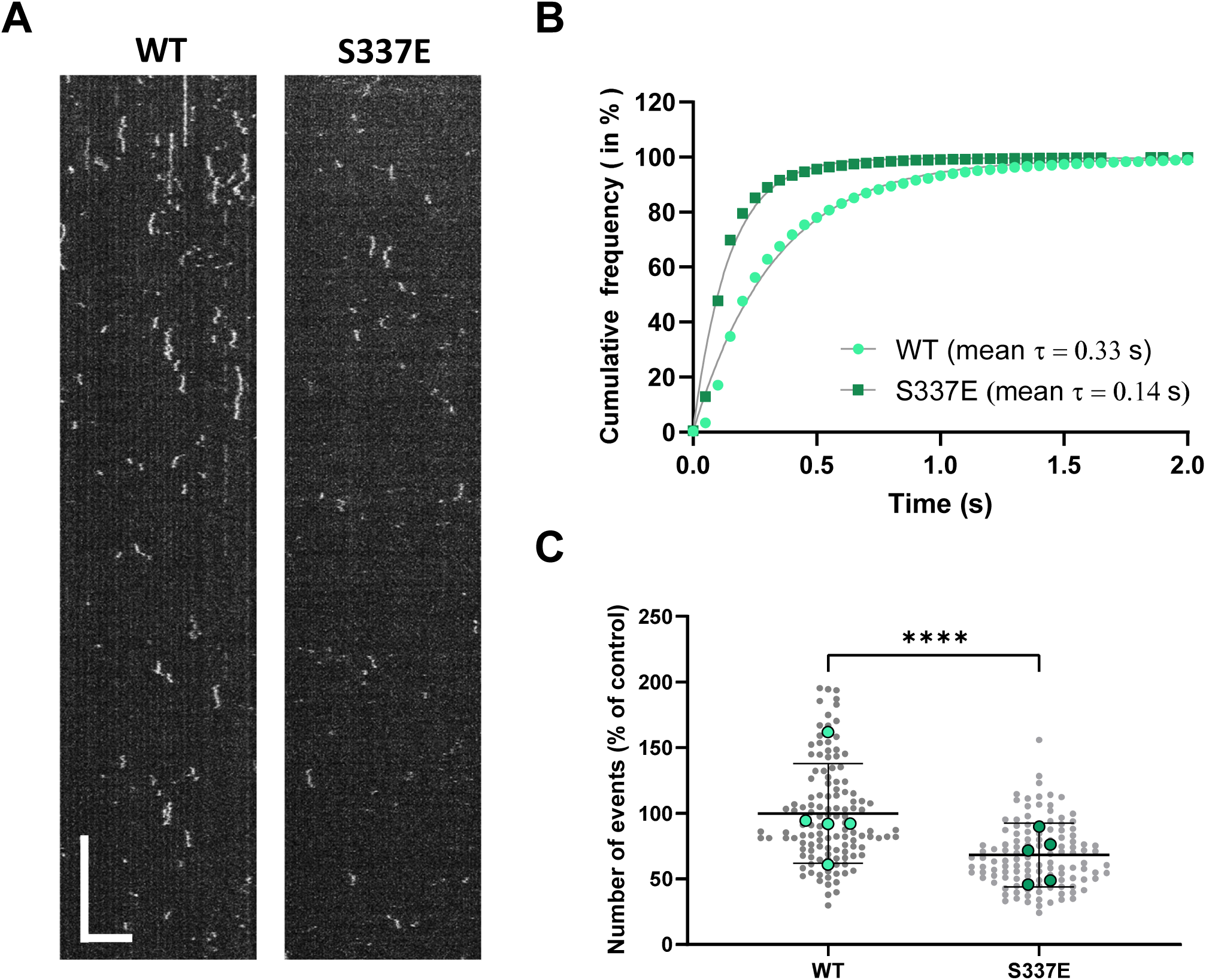
The VASH1 C-terminal phosphomimic mutant S337E shows decreased binding to taxol-stabilized microtubules. Comparison of the interaction of the WT and mutant enzyme with microtubules by single molecule TIRF microscopy in the same experimental conditions as in Figure 5. **(A)**. Representative kymographs. Scale bars: horizontal, 5 μm; vertical, 5 s. **(B-C)** Analysis of the binding characteristics of the two VASH-SVBP enzymes from 5 independent experiments (done with 4 and 3 different protein preparations, respectively for WT and S337E). **(B)** Cumulative frequency of the residence times measured in TIRF movies following enzyme addition. The mean residence time (τ) is obtained by fitting the curve with a mono-exponential function. **(C)** Analysis of binding frequency. Each point represents a microtubule. Around 20 microtubules were analyzed per experiment, 5 independent experiments. Data are represented as the median with the interquartile range. Statistical significance was determined using Kruskal-Wallis test. ****p < 0.0001.

**Figure S6.**
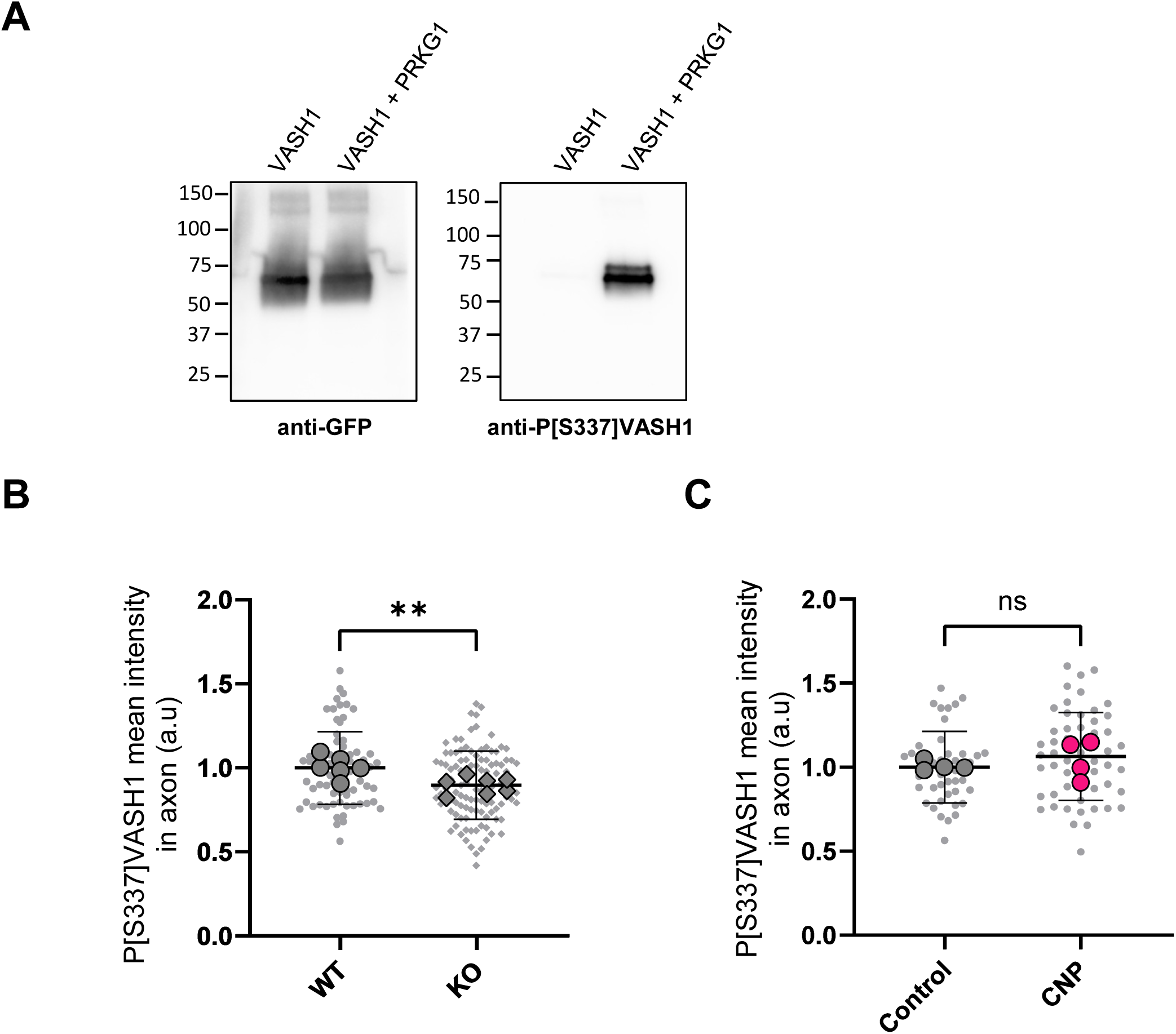
Characterization of anti-P[S337]VASH1 antibody and comparison of immunofluorescence signals for P[S337]VASH1 in the axons of WT and *Vash1* KO neurons. (**A**) Recombinant sfGFP-VASH1-SVBP (2 µM) was incubated without or with PRKG1α (10 nM) and ATP for 1 h and analyzed by immunoblot using anti-GFP and anti-P[S337]VASH1 antibody. GFP content confirms similar loading of the samples. (**B**) Quantification of P[S337]VASH1 in axons of WT and SVBP-KO neurons. n/N: 73/6 (WT), 107/7 (KO) with n indicating the number of axons analyzed and N the number of embryos. (**C**) Quantification of P[S337]VASH1 in axons of WT neurons subjected or not to CNP. n/N: 45/4 (WT), 53/4 (KO) with n indicating the number of axons analyzed and N the number of embryos.

**Table S1.** Peptides identified by MS-based proteomics in the C-terminal region of VASH1.

| peptide sequence | ptm protein positions | ptm sites confidence | psm score | accession | start | end | SC R1-Trp | SC R2-Trp | SC R3-Trp | SC R1-LysC | SC R2-LysC | SC R3-LysC |
| --- | --- | --- | --- | --- | --- | --- | --- | --- | --- | --- | --- | --- |
| IGKGTGPPSPTKDR |  |  | 140,37 | sfGFP-hVash1-myc-His | 554 | 567 | 6 | 4 |  |  |  |  |
| GKGTGPPSPTKDR |  |  | 33,25 | sfGFP-hVash1-myc-His | 555 | 567 |  | 1 |  |  |  |  |
| GTGPPSPTK |  |  | 56,97 | sfGFP-hVash1-myc-His | 557 | 565 |  | 1 | 1 |  |  | 2 |
| GTGPPSPTKDR |  |  | 35,82 | sfGFP-hVash1-myc-His | 557 | 567 |  |  |  |  |  | 1 |
| GTGPPSPTKDRK |  |  | 91,66 | sfGFP-hVash1-myc-His | 557 | 568 |  |  |  | 10 | 5 | 1 |
| GPPSPTKDRK |  |  | 47,57 | sfGFP-hVash1-myc-His | 559 | 568 |  |  |  | 1 | 1 |  |
| KDVSSPQR |  |  | 52,69 | sfGFP-hVash1-myc-His | 569 | 576 |  |  |  |  |  | 1 |
| KDVSSPQRAQSSPH |  |  | 35,09 | sfGFP-hVash1-myc-His | 569 | 582 |  |  |  | 1 |  |  |
| KDVSSPQRAQSSPHRR |  |  | 64,92 | sfGFP-hVash1-myc-His | 569 | 584 |  |  |  | 2 | 4 |  |
| DVSSPQR |  |  | 29,38 | sfGFP-hVash1-myc-His | 570 | 576 |  |  | 1 |  |  | 1 |
| DVSSPQRAQSSPHRR |  |  | 40,92 | sfGFP-hVash1-myc-His | 570 | 584 |  |  |  | 2 | 5 |  |
| SSPQRAQSSPHRR |  |  | 47,96 | sfGFP-hVash1-myc-His | 572 | 584 |  |  |  |  | 1 |  |
| TSEPKAMPDLNGYQIR | Oxidation (M604) | Oxidation (M7) = 1.0 | 88,51 | sfGFP-hVash1-myc-His | 598 | 613 |  | 1 |  |  |  |  |
| TSEPKAMPDLNGYQIRVTRTRPLEQK | Oxidation (M604) | Oxidation (M7) = 1.0 | 40,73 | sfGFP-hVash1-myc-His | 598 | 623 |  |  |  |  | 1 |  |
| AMPDLNGYQIR |  |  | 84,98 | sfGFP-hVash1-myc-His | 603 | 613 | 1 | 2 | 1 | 1 | 1 | 2 |
| AMPDLNGYQIR | Oxidation (M604) | Oxidation (M2) = 1.0 | 80,86 | sfGFP-hVash1-myc-His | 603 | 613 | 2 | 1 | 2 | 1 | 1 | 1 |
| AMPDLNGYQIRVT |  |  | 85,88 | sfGFP-hVash1-myc-His | 603 | 615 |  |  |  | 1 |  |  |
| AMPDLNGYQIRVT | Oxidation (M604) | Oxidation (M2) = 1.0 | 93,38 | sfGFP-hVash1-myc-His | 603 | 615 |  |  |  | 1 |  |  |
| AMPDLNGYQIRVTR |  |  | 74,1 | sfGFP-hVash1-myc-His | 603 | 616 |  |  |  | 2 | 2 |  |
| AMPDLNGYQIRVTR | Oxidation (M604) | Oxidation (M2) = 1.0 | 73,63 | sfGFP-hVash1-myc-His | 603 | 616 |  |  |  | 2 | 2 | 1 |
| AMPDLNGYQIRVTRTRPLEQK | Oxidation (M604) | Oxidation (M2) = 1.0 | 24,53 | sfGFP-hVash1-myc-His | 603 | 623 |  |  |  |  | 1 |  |
| MPDLNGYQIR | Oxidation (M604) | Oxidation (M1) = 1.0 | 40,49 | sfGFP-hVash1-myc-His | 604 | 613 |  | 1 |  |  |  |  |
| PDLNGYQIR |  |  | 95,37 | sfGFP-hVash1-myc-His | 605 | 613 |  | 1 |  |  |  |  |

**Table S2.**
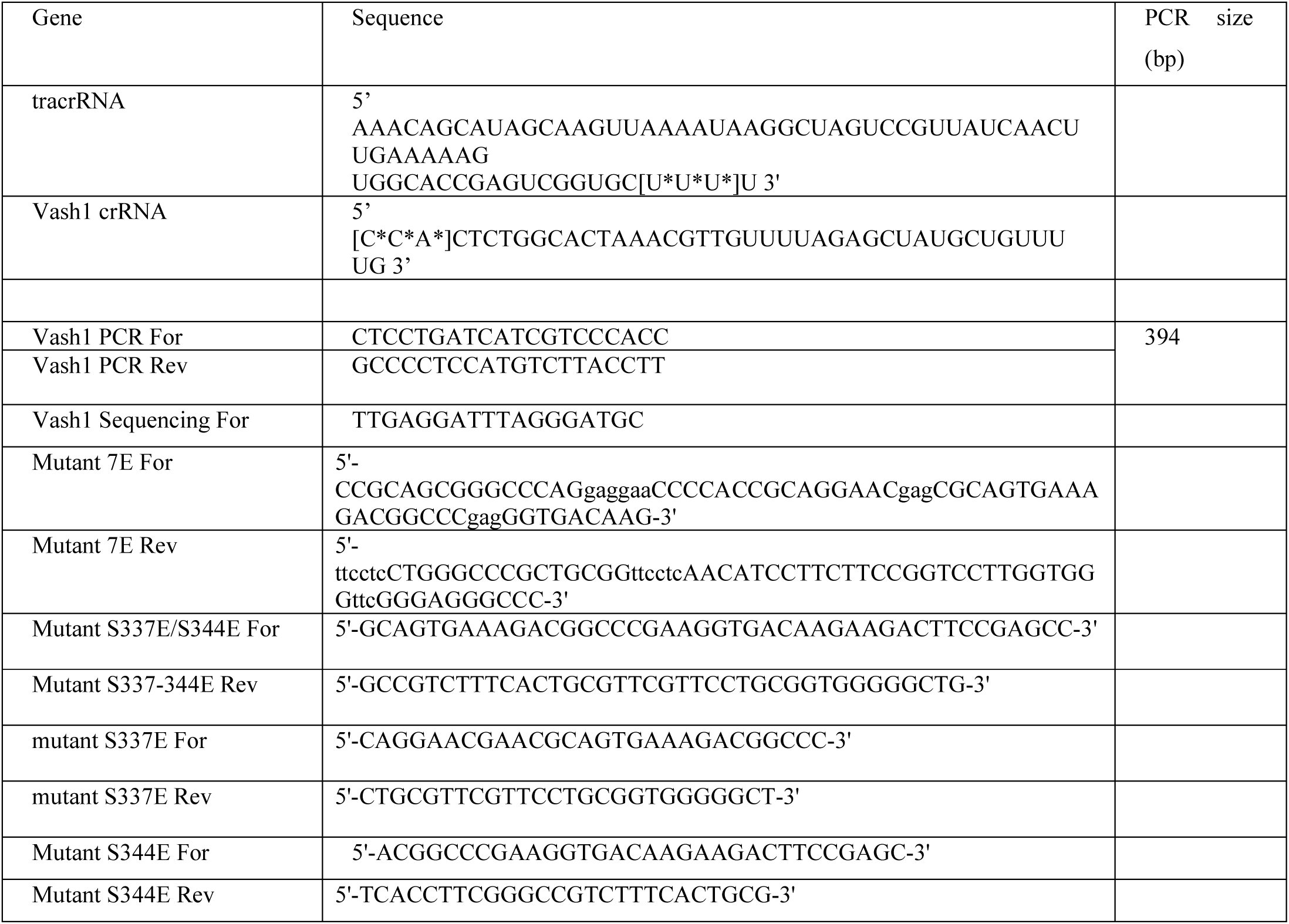
Oligonucleotides used in the study. Note: * are for phosphothiorate and [] for 2’-O-methyl chemical modifications.

## REFERENCES

Ahmad, F.J., T.P. Pienkowski, and P.W. Baas. 1993. Regional differences in microtubule dynamics in the axon. J Neurosci. 13:856–866.

Aillaud, C., C. Bosc, L. Peris, A. Bosson, P. Heemeryck, J. Van Dijk, J. Le Friec, B. Boulan, F. Vossier, L.E. Sanman, S. Syed, N. Amara, Y. Coute, L. Lafanechere, E. Denarier, C. Delphin, L. Pelletier, S. Humbert, M. Bogyo, A. Andrieux, K. Rogowski, and M.J. Moutin. 2017. Vasohibins/SVBP are tubulin carboxypeptidases (TCPs) that regulate neuron differentiation. Science. 358:1448–1453.

Aillaud, C., C. Bosc, Y. Saoudi, E. Denarier, L. Peris, L. Sago, N. Taulet, A. Cieren, O. Tort, M.M. Magiera, C. Janke, V. Redeker, A. Andrieux, and M.J. Moutin. 2016. Evidence for new C-terminally truncated variants of alpha- and beta-tubulins. Mol Biol Cell. 27:640–653.

Akiyama, H., T. Fukuda, T. Tojima, V.O. Nikolaev, and H. Kamiguchi. 2016. Cyclic Nucleotide Control of Microtubule Dynamics for Axon Guidance. J Neurosci. 36:5636–5649.

Baas, P.W., and M.M. Black. 1990. Individual microtubules in the axon consist of domains that differ in both composition and stability. J Cell Biol. 111:495–509.

Balmes, A., H. Schmidt, S. Peters, S. Kenet, A. Botezatu, L. Wen, A. Bottcher, P.M. Benz, R. Feil, and T.E. Schaffer. 2026. CNP-induced cGMP signaling reduces growth cone stiffness and Ca(2+) levels in embryonic DRG neurons. Front Mol Neurosci. 19:1769175.

Banko, M.R., J.J. Allen, B.E. Schaffer, E.W. Wilker, P. Tsou, J.L. White, J. Villen, B. Wang, S.R. Kim, K. Sakamoto, S.P. Gygi, L.C. Cantley, M.B. Yaffe, K.M. Shokat, and A. Brunet. 2011. Chemical genetic screen for AMPKalpha2 substrates uncovers a network of proteins involved in mitosis. Mol Cell. 44:878–892.

Bouquet, C., and F. Nothias. 2007. Molecular mechanisms of axonal growth. Adv Exp Med Biol. 621:1–16.

Bouyssie, D., A.M. Hesse, E. Mouton-Barbosa, M. Rompais, C. Macron, C. Carapito, A. Gonzalez de Peredo, Y. Coute, V. Dupierris, A. Burel, J.P. Menetrey, A. Kalaitzakis, J. Poisat, A. Romdhani, O. Burlet-Schiltz, S. Cianferani, J. Garin, and C. Bruley. 2020. Proline: an efficient and user-friendly software suite for large-scale proteomics. Bioinformatics. 36:3148–3155.

Brown, A., Y. Li, T. Slaughter, and M.M. Black. 1993. Composite microtubules of the axon: quantitative analysis of tyrosinated and acetylated tubulin along individual axonal microtubules. J Cell Sci. 104 (Pt 2):339–352.

Casabona, M.G., Y. Vandenbrouck, I. Attree, and Y. Coute. 2013. Proteomic characterization of Pseudomonas aeruginosa PAO1 inner membrane. Proteomics. 13:2419–2423.

Chen, C.Y., A.K. Salomon, M.A. Caporizzo, S. Curry, N.A. Kelly, K. Bedi, A.I. Bogush, E. Kramer, S. Schlossarek, P. Janiak, M.J. Moutin, L. Carrier, K.B. Margulies, and B.L. Prosser. 2020. Depletion of Vasohibin 1 Speeds Contraction and Relaxation in Failing Human Cardiomyocytes. Circ Res. 127:e14–e27.

Conde, C., and A. Caceres. 2009. Microtubule assembly, organization and dynamics in axons and dendrites. Nat Rev Neurosci. 10:319–332.

Coute, Y., C. Bruley, and T. Burger. 2020. Beyond Target-Decoy Competition: Stable Validation of Peptide and Protein Identifications in Mass Spectrometry-Based Discovery Proteomics. Anal Chem. 92:14898–14906.

Dent, E.W., and F.B. Gertler. 2003. Cytoskeletal dynamics and transport in growth cone motility and axon guidance. Neuron. 40:209–227.

Dent, E.W., and K. Kalil. 2001. Axon branching requires interactions between dynamic microtubules and actin filaments. J Neurosci. 21:9757–9769.

Dumoulin, A., A. Dagane, G. Dittmar, and F.G. Rathjen. 2018a. S-palmitoylation Is Required for the Control of Growth Cone Morphology of DRG Neurons by CNP-Induced cGMP Signaling. Front Mol Neurosci. 11:345.

Dumoulin, A., H. Schmidt, and F.G. Rathjen. 2021. Sensory Neurons: The Formation of T-Shaped Branches Is Dependent on a cGMP-Dependent Signaling Cascade. Neuroscientist. 27:47–57.

Dumoulin, A., G. Ter-Avetisyan, H. Schmidt, and F.G. Rathjen. 2018b. Molecular Analysis of Sensory Axon Branching Unraveled a cGMP-Dependent Signaling Cascade. Int J Mol Sci. 19.

Erck, C., L. Peris, A. Andrieux, C. Meissirel, A.D. Gruber, M. Vernet, A. Schweitzer, Y. Saoudi, H. Pointu, C. Bosc, P.A. Salin, D. Job, and J. Wehland. 2005. A vital role of tubulin-tyrosine-ligase for neuronal organization. Proc Natl Acad Sci U S A. 102:7853–7858.

Ersfeld, K., J. Wehland, U. Plessmann, H. Dodemont, V. Gerke, and K. Weber. 1993. Characterization of the tubulin-tyrosine ligase. J Cell Biol. 120:725–732.

Haeussler, M., K. Schonig, H. Eckert, A. Eschstruth, J. Mianne, J.B. Renaud, S. Schneider-Maunoury, A. Shkumatava, L. Teboul, J. Kent, J.S. Joly, and J.P. Concordet. 2016. Evaluation of off-target and on-target scoring algorithms and integration into the guide RNA selection tool CRISPOR. Genome Biol. 17:148.

Iqbal, Z., H. Tawamie, W. Ba, A. Reis, B.A. Halak, H. Sticht, S. Uebe, N.N. Kasri, S. Riazuddin, H. van Bokhoven, and R. Abou Jamra. 2019. Loss of function of SVBP leads to autosomal recessive intellectual disability, microcephaly, ataxia, and hypotonia. Genet Med. 21:1790–1796.

Janke, C., and M.M. Magiera. 2020. The tubulin code and its role in controlling microtubule properties and functions. Nat Rev Mol Cell Biol. 21:307–326.

Kalil, K., and E.W. Dent. 2005. Touch and go: guidance cues signal to the growth cone cytoskeleton. Curr Opin Neurobiol. 15:521–526.

Kim, W.Y., F.Q. Zhou, J. Zhou, Y. Yokota, Y.M. Wang, T. Yoshimura, K. Kaibuchi, J.R. Woodgett, E.S. Anton, and W.D. Snider. 2006. Essential roles for GSK-3s and GSK-3-primed substrates in neurotrophin-induced and hippocampal axon growth. Neuron. 52:981–996.

Kishimoto, I., T. Tokudome, T. Horio, T. Soeki, H. Chusho, K. Nakao, and K. Kangawa. 2008. C-type natriuretic peptide is a Schwann cell-derived factor for development and function of sensory neurones. J Neuroendocrinol. 20:1213–1223.

Landskron, L., J. Bak, A. Adamopoulos, K. Kaplani, M. Moraiti, L.G. van den Hengel, J.Y. Song, O.B. Bleijerveld, J. Nieuwenhuis, T. Heidebrecht, L. Henneman, M.J. Moutin, M. Barisic, S. Taraviras, A. Perrakis, and T.R. Brummelkamp. 2022. Posttranslational modification of microtubules by the MATCAP detyrosinase. Science. 376:eabn6020.

Liz, M.A., F.M. Mar, T.E. Santos, H.I. Pimentel, A.M. Marques, M.M. Morgado, S. Vieira, V.F. Sousa, H. Pemble, T. Wittmann, C. Sutherland, J.R. Woodgett, and M.M. Sousa. 2014. Neuronal deletion of GSK3beta increases microtubule speed in the growth cone and enhances axon regeneration via CRMP-2 and independently of MAP1B and CLASP2. BMC Biol. 12:47.

Lowery, L.A., and D. Van Vactor. 2009. The trip of the tip: understanding the growth cone machinery. Nat Rev Mol Cell Biol. 10:332–343.

Marcos, S., J. Moreau, S. Backer, D. Job, A. Andrieux, and E. Bloch-Gallego. 2009. Tubulin tyrosination is required for the proper organization and pathfinding of the growth cone. PLoS One. 4:e5405.

Markossian, S., R. Guyot, S. Richard, M. Teixeira, N. Aguilera, M. Bouchet, M. Plateroti, W. Guan, K. Gauthier, D. Aubert, and F. Flamant. 2018. CRISPR/Cas9 Editing of the Mouse Thra Gene Produces Models with Variable Resistance to Thyroid Hormone. Thyroid. 28:139–150.

Meyer-Jens M, S.C., Pietsch N., Ramirez-Rios S., Herrera-Rivero M., Krämer E., Braren I., Nikolaev V., Frye M., Könemann S., Schlossarek S., Moutin M.-J., Carrier L.. 2026. Neprilysin inhibition reduces microtubule detyrosination in cardiomyocytes through a cGMP-PRKG1-VASH1 axis. bioRxiv.

Moutin, M.J., C. Bosc, L. Peris, and A. Andrieux. 2021. Tubulin post-translational modifications control neuronal development and functions. Dev Neurobiol. 81:253–272.

Nascimento, A.I., F.M. Mar, and M.M. Sousa. 2018. The intriguing nature of dorsal root ganglion neurons: Linking structure with polarity and function. Prog Neurobiol. 168:86–103.

Nicot, S., G. Gillard, H. Impheng, E. Joachimiak, S. Urbach, K. Mochizuki, D. Wloga, F. Juge, and K. Rogowski. 2023. A family of carboxypeptidases catalyzing alpha- and beta-tubulin tail processing and deglutamylation. Sci Adv. 9:eadi7838.

Nieuwenhuis, J., A. Adamopoulos, O.B. Bleijerveld, A. Mazouzi, E. Stickel, P. Celie, M. Altelaar, P. Knipscheer, A. Perrakis, V.A. Blomen, and T.R. Brummelkamp. 2017. Vasohibins encode tubulin detyrosinating activity. Science. 358:1453–1456.

Pagnamenta, A.T., P. Heemeryck, H.C. Martin, C. Bosc, L. Peris, I. Uszynski, S. Gory-Faure, S. Couly, C. Deshpande, A. Siddiqui, A.A. Elmonairy, W.G.S. Consortium, C. Genomics England Research, S. Jayawant, S. Murthy, I. Walker, L. Loong, P. Bauer, F. Vossier, E. Denarier, T. Maurice, E.L. Barbier, J.C. Deloulme, J.C. Taylor, E.M. Blair, A. Andrieux, and M.J. Moutin. 2019. Defective tubulin detyrosination causes structural brain abnormalities with cognitive deficiency in humans and mice. Hum Mol Genet. 28:3391–3405.

Perez-Riverol, Y., J. Bai, C. Bandla, D. Garcia-Seisdedos, S. Hewapathirana, S. Kamatchinathan, D.J. Kundu, A. Prakash, A. Frericks-Zipper, M. Eisenacher, M. Walzer, S. Wang, A. Brazma, and J.A. Vizcaino. 2022. The PRIDE database resources in 2022: a hub for mass spectrometry-based proteomics evidences. Nucleic Acids Res. 50:D543–D552.

Peris, L., M. Wagenbach, L. Lafanechere, J. Brocard, A.T. Moore, F. Kozielski, D. Job, L. Wordeman, and A. Andrieux. 2009. Motor-dependent microtubule disassembly driven by tubulin tyrosination. J Cell Biol. 185:1159–1166.

Ramirez-Rios, S., S.R. Choi, C. Sanyal, T.B. Blum, C. Bosc, F. Krichen, E. Denarier, J.M. Soleilhac, B. Blot, C. Janke, V. Stoppin-Mellet, M.M. Magiera, I. Arnal, M.O. Steinmetz, and M.J. Moutin. 2023. VASH1-SVBP and VASH2-SVBP generate different detyrosination profiles on microtubules. J Cell Biol. 222.

Ramirez-Rios, S., L. Serre, V. Stoppin-Mellet, E. Prezel, A. Vinit, E. Courriol, A. Fourest-Lieuvin, J. Delaroche, E. Denarier, and I. Arnal. 2017. A TIRF microscopy assay to decode how tau regulates EB’s tracking at microtubule ends. Methods Cell Biol. 141:179–197.

Sanyal, C., N. Pietsch, S. Ramirez Rios, L. Peris, L. Carrier, and M.J. Moutin. 2021. The detyrosination/re-tyrosination cycle of tubulin and its role and dysfunction in neurons and cardiomyocytes. Semin Cell Dev Biol.

Schmidt, H., A. Stonkute, R. Juttner, D. Koesling, A. Friebe, and F.G. Rathjen. 2009. C-type natriuretic peptide (CNP) is a bifurcation factor for sensory neurons. Proc Natl Acad Sci U S A. 106:16847–16852.

Schmidt, H., M. Werner, P.A. Heppenstall, M. Henning, M.I. More, S. Kuhbandner, G.R. Lewin, F. Hofmann, R. Feil, and F.G. Rathjen. 2002. cGMP-mediated signaling via cGKIalpha is required for the guidance and connectivity of sensory axons. J Cell Biol. 159:489–498.

Souphron, J., S. Bodakuntla, A.S. Jijumon, G. Lakisic, A.M. Gautreau, C. Janke, and M.M. Magiera. 2019. Purification of tubulin with controlled post-translational modifications by polymerization-depolymerization cycles. Nat Protoc. 14:1634–1660.

Wehland, J., and K. Weber. 1987. Tubulin-tyrosine ligase has a binding site on beta-tubulin: a two-domain structure of the enzyme. J Cell Biol. 104:1059–1067.

Wood, J.S., X. Yan, M. Mendelow, J.D. Corbin, S.H. Francis, and D.S. Lawrence. 1996. Precision substrate targeting of protein kinases. The cGMP- and cAMP-dependent protein kinases. J Biol Chem. 271:174–179.

Xia, C., M. Nguyen, A.K. Garrison, Z. Zhao, Z. Wang, C. Sutherland, and L. Ma. 2013. CNP/cGMP signaling regulates axon branching and growth by modulating microtubule polymerization. Dev Neurobiol. 73:673–687.

Zhao, Z., and L. Ma. 2009. Regulation of axonal development by natriuretic peptide hormones. Proc Natl Acad Sci U S A. 106:18016–18021.

